# A plasma proteomic atlas of rare genetic neurodevelopmental disorders reveals convergent molecular signatures, neurodegenerative overlap, and a cross-disorder severity index

**DOI:** 10.64898/2026.11.16.767059

**Authors:** Ricardo N Ramirez, Megan A Aumann, Lindsay Wieczorek, Sasha Elmizadeh, Albert Misko, Yael Weiss, Stephanie Szobota, Berit Powers, Shruti Sasaki, Marco Rizzo, Smitha Jagadish, Mariana Laranjo, Nicole Davis, Jérémy Manry, Marie-Elodie Cattin-Messaoudi, Joerg Hipp, Brett Younginger, Rachael Hawtin, Nichol Miller, Kevin Wright, Grace Viggiano, Sarah Poliquin, Taylor Morris, Zollie Yavarow, Nicholas Aguilar, Ananya Terala, William Keener, Kristin Hatcher, Christopher Ricupero, Jennifer Bain, Elizabeth Berry Kravitz, Jacqueline Harris, Natasha N Ludwig, Angela Duong, Wayne W Poon, Terry Jo V Bichell, Anna C Pfalzer

## Abstract

Rare genetic neurodevelopmental disorders (RGNDs) are individually too small for well-powered biomarker studies. We profiled over 10,000 plasma proteins across 27 RGNDs and unaffected family controls, alongside standardized adaptive function measures, to identify shared molecular features. Cross-disorder analysis revealed two proteomic clusters distinguished by cytoskeletal and vesicle-trafficking versus metabolic and hepatic proteins. We identified recurrent protein changes and a shared 32-protein epilepsy signature largely undetectable within individual disorders. RGND proteomic signatures also overlapped extensively with those of adult neurodegenerative diseases, including Alzheimer’s, Parkinson’s, frontotemporal dementia, and amyotrophic lateral sclerosis, often in opposing directions. A 186-protein plasma index tracked communication ability in disorders excluded from model training and among individuals with the same diagnosis, independently of age. Twenty-four prioritized proteins have existing drug interactions, including targets of approved agents. These findings demonstrate that cross-disorder proteomics reveals otherwise inaccessible molecular signatures, clinically relevant biomarkers, and potentially actionable biology across genetically distinct RGNDs.

## MAIN

A rare disease is defined as a condition that affects less than 200,000 individuals in the US^1^. There are an estimated 10,000+ rare diseases that affect more than 30 million people – approximately one out of every 10 people – in the U.S.^2^, and about half of these people are children3. Despite over 700 clinical trial grants awarded by the FDA Office of Orphan Products Development, thousands of rare diseases remain without approved treatments. Among the most severely affected are rare genetic neurodevelopmental disorders (RGNDs), where a single-gene lesion disrupts brain development and objective measures of disease state are largely unknown.

A major obstacle for establishing rare disease signatures is cohort size. Each RGND is individually too rare to support a well-powered biomarker study, and each is conventionally studied on its own, so any shared biology across disorders remains invisible by construction. Drug development is further challenged by phenotypic heterogeneity, limited understanding of natural history, and the global dispersion of patients, impacting biomarker discovery. Studying these disorders together would address the sample-size problem directly, but requires samples and phenotypes collected using a shared protocol. The patient-advocacy governed non-profit Consortium for Outcome Measures and Biomarkers for Neurodevelopmental Disorders (COMBINEDBrain) established a biorepository to facilitate biospecimen collection and phenotypic data aggregation across RGNDs^4^. By linking samples with a uniform dataset encompassing standardized clinical, neuropsychological, and demographic data, a major consortium aim was to increase statistical power to enable biomarker discovery across RGNDs.

Plasma is a well-suited biological specimen to address this question: it is accessible in pediatric populations whereas cerebrospinal fluid and imaging are difficult to obtain longitudinally; plasma carries proteins reflecting immune, metabolic, and neuronal processes, enabling multiplexing. This study describes a plasma proteomic resource developed from sample and data aggregation across 27 RGNDs collected through COMBINEDBrain, 26 of which were carried forward to differential abundance analysis. This resource defined shared and disorder-specific proteomic signatures, identified convergent pathways and druggable targets, uncovered overlap with neurodegenerative disease, and established a multi-protein severity index tied to developmental outcomes. For many of the disorders, this represents the first systematic investigation of plasma-based biomarkers nominating RGND-specific biomarkers for testing efficacy of therapeutic candidates. The RGND proteomics data and analytical pipelines serve as a community resource to accelerate biomarker prioritization, drug target nomination, and clinical trial design across the rare neurodevelopmental disease landscape.

## RESULTS

### Plasma proteins reflect causal mutations and shared biology across disorders

Cognate proteins, the proteins encoded by genes directly implicated in each disorder, provide a direct test of whether plasma proteomics can capture the molecular effects of rare genetic neurodevelopmental disorders (RGNDs). Examination of the Human Protein Atlas revealed that almost all RGND cognate proteins were peripherally expressed; NR2F1 expression has yet to be determined. Using the SomaScan® 11K Assay platform, we profiled 10,738 plasma proteins in 522 individuals across 27 RGNDs and 172 unaffected sibling or family controls (Figure 1, Table 1).

**Figure 1.**
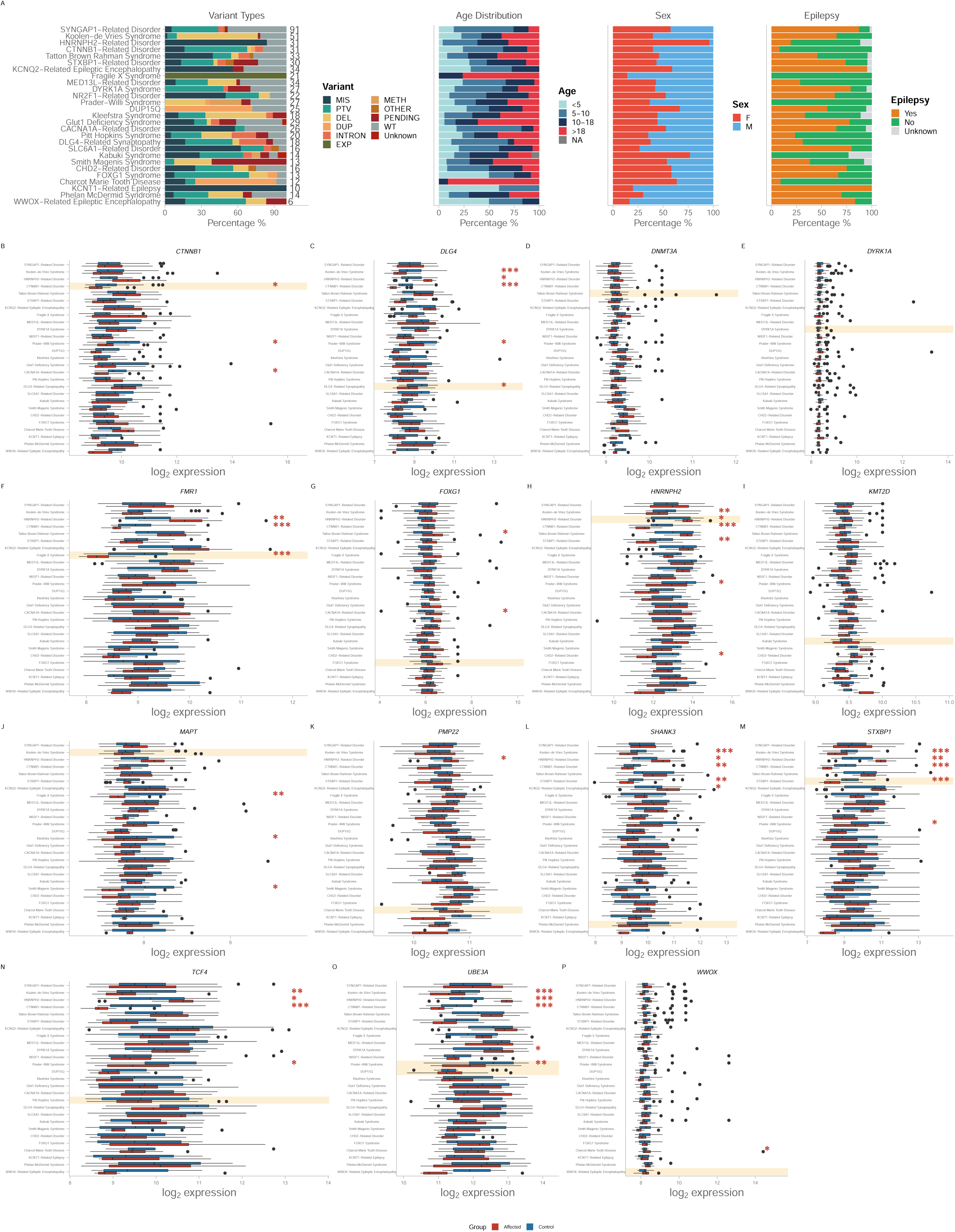
Cohort composition and plasma expression of cognate disease proteins across the 27 recruited rare genetic neurodevelopmental disorders. (A) Cohort demographics and clinical characteristics. Stacked bar charts display the distribution of variant types (all samples, including controls), age groups (<5, 5-10, 10-18, >18 years), sex (female, male), and epilepsy status (yes, no, unknown) across all 27 recruited rare neurodevelopmental disorders, ordered by decreasing affected sample size; 26 of these met the eight-participant threshold for the differential abundance analyses in Figures 2 to 6. Total sample counts are annotated to the right of each bar. Controls (n = 172) and affected individuals (n = 522) were profiled using SomaScan v5.0 (EDTA plasma, ANML normalized; 10,738 proteins). Variant types include missense (MIS), protein-truncating variants (PTV), deletions (DEL), duplications (DUP), intronic (INTRON), repeat expansions (EXP), methylation (METH), and other/pending categories. (B-P) Cognate protein expression across the recruited disorders. Each panel shows log2-transformed plasma abundance (RFU) of a single disorder-causal protein, displayed as side-by-side boxplots for affected (red) and control (blue) groups. The cognate disorder for each protein is highlighted with a yellow background. Significance is from per-disorder linear models (log2(RFU) ∼ Group + Sex + Age_scaled, limma with empirical Bayes moderation), reported as Benjamini-Hochberg adjusted FDR < 0.10 with |log2 fold-change| > 0.2, the criterion used throughout. Panels: (B) CTNNB1/beta-catenin, (C) DLG4/PSD-95, (D) DNMT3A, (E) DYRK1A, (F) FMR1/FMRP, (G) FOXG1, (H) HNRNPH2, (I) KMT2D, (J) MAPT/tau (cognate disorder: KANSL1/Koolen-de Vries, sharing the 17q21.31 locus), (K) PMP22, (L) SHANK3, (M) STXBP1/Munc18-1, (N) TCF4, (O) UBE3A/E6AP (cognate disorders: PAT15Q and DUP15Q, sharing the 15q11-q13 region), (P) WWOX. Panel P is shown for completeness: WWOX-Related Epileptic Encephalopathy contributed six affected participants, below the eight-participant threshold, and was not carried into the differential abundance analyses.

**Table 1.**
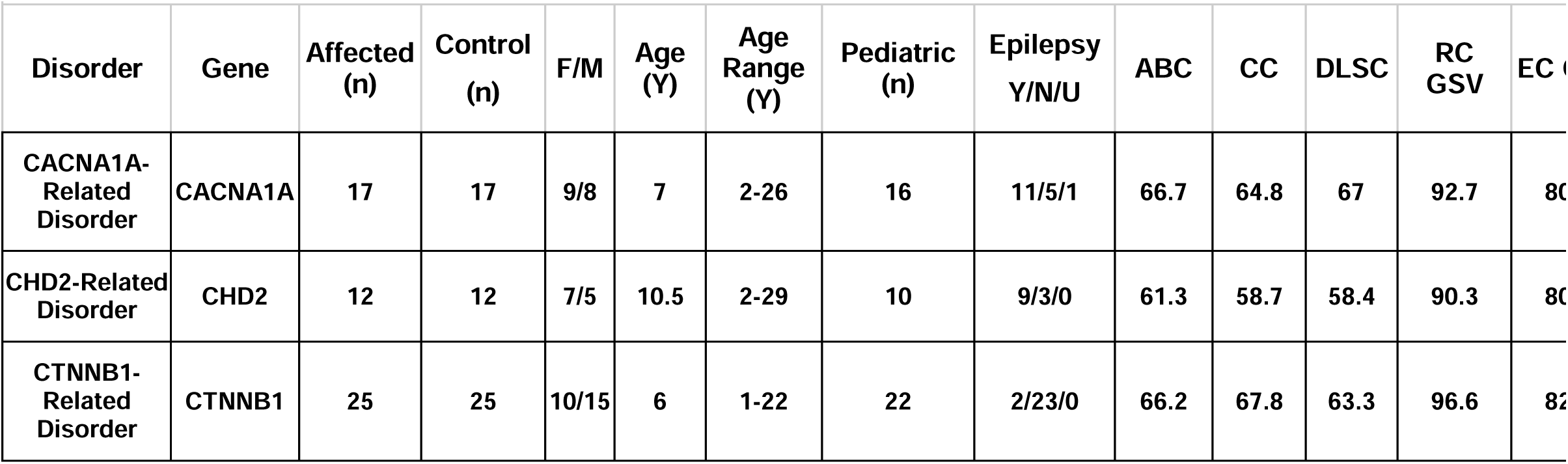
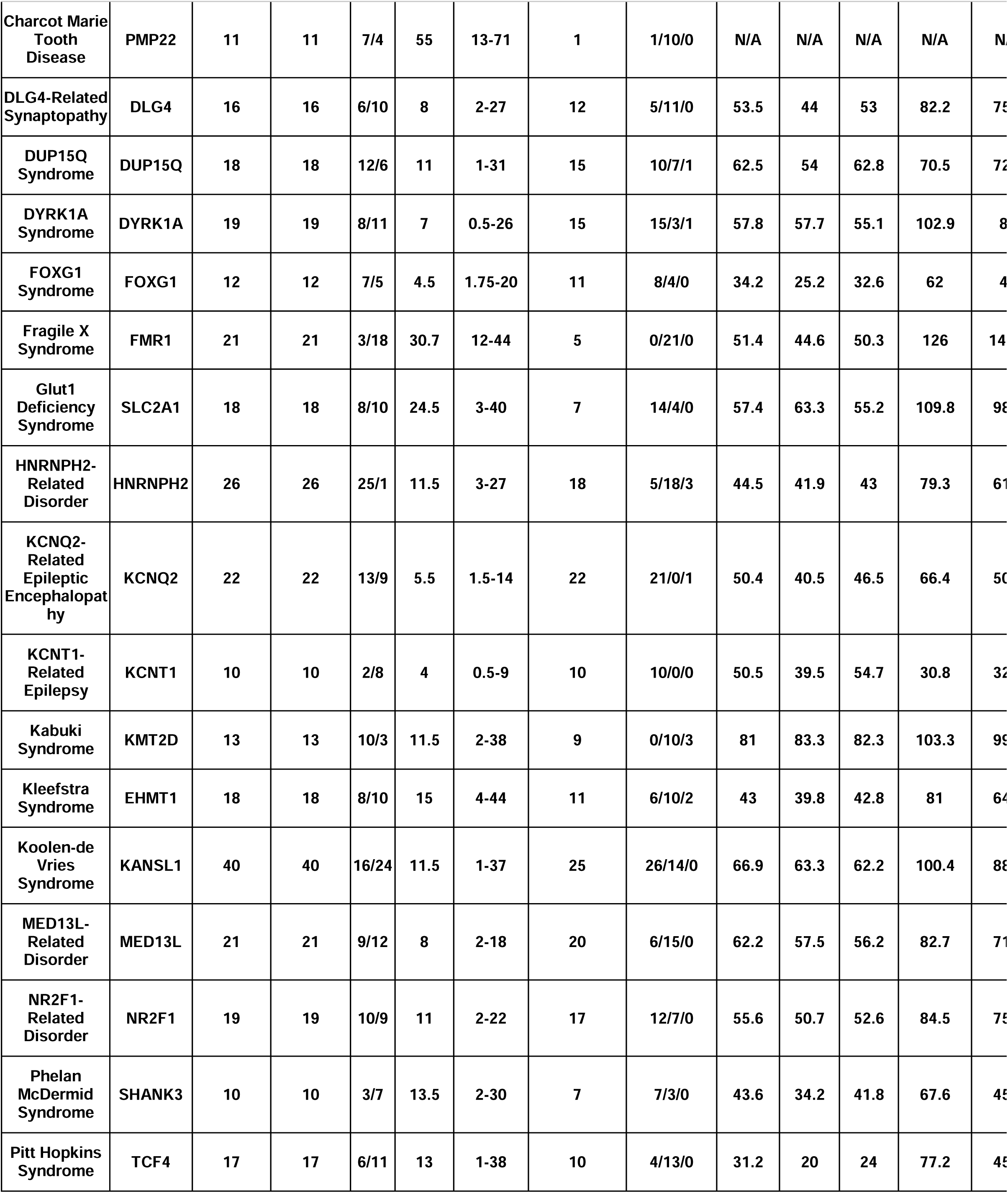
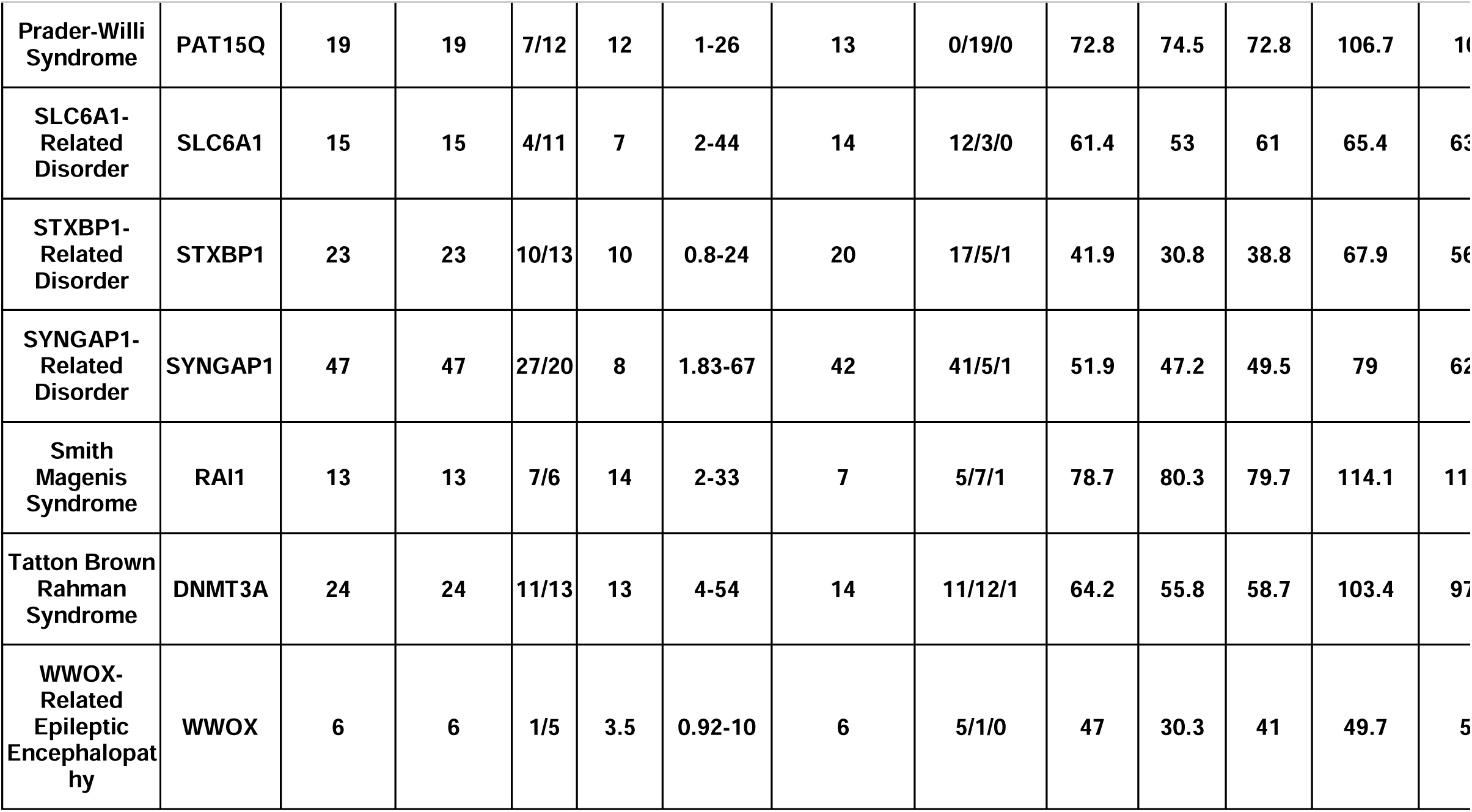
Clinical Characteristics of Rare Genetic Neurological Disease (RGND) cohort. The number of affected and unaffected (Control) participants with twenty-seven different genetic diseases are shown with mean and range of age, male:female (M/F) ratio and number of participants under the age of 18 years (Pediatrics). Epilepsy status for all participants was also evaluated using: Yes(Y), No(N), Unknown (U). Affected participant Vineland aberrant behavior (ABC), communication (CC) and daily living skills composite (DLSC), and growth-scale values for receptive communication (RCGSV) and expressive communication (ECGSV) are shown.

Plasma levels of several cognate proteins reflected the underlying genetic changes. βeta-catenin (CTNNB1), PSD-95 (DLG4), FMR1, STXBP1 and UBE3A were significantly reduced in their respective haploinsufficiency disorders (Figure 1B, 1C, 1F, 1M, 1O). In contrast, HNRNPH2 was elevated in HNRNPH2-Related Disorder (Figure 1H), potentially reflecting altered localization of the mutant protein. No significant changes were detected for DNMT3A, DYRK1A, FOXG1, KMT2D, MAPT, PMP22 or SHANK3 in their corresponding disorders (Figure 1D, 1E, 1G, 1I, 1J, 1K, 1L). Together, these findings suggest that plasma cognate protein levels can directly reflect disorders caused by reduced protein abundance but may be less informative when disease results from altered protein function, such as dominant-negative or gain-of-function variants.

Importantly, changes in these proteins were not limited to their corresponding disorders. For example, βeta-catenin was reduced not only in CTNNB1-Related Disorder, but also in Prader-Willi Syndrome and CACNA1A-Related Disorder (Figure 1B). Similarly, the synaptic proteins PSD-95 and STXBP1 were altered across several genetically distinct disorders, including Koolen-de Vries Syndrome, HNRNPH2-Related Disorder, CTNNB1-Related Disorder and Prader-Willi Syndrome (Figure 1C, 1M). TCF4 and UBE3A were also altered across multiple disorders (Figure 1N, 1O). Therefore, plasma proteins capture both direct effects of disease-causing genetic variants and molecular changes shared across genetically distinct disorders, revealing convergence on common synaptic and transcriptional pathways.

### Rare neurodevelopmental disorders show diverse plasma proteomic signatures

Beyond changes in cognate proteins, we examined the broader plasma proteome to determine whether each RGND was associated with a detectable proteomic signature. Among the 16 disorders in which recruitment site could be accounted for, 15 showed significant proteomic changes compared to control, with 2,484 differentially abundant proteins identified across disorders (Figure 2A, Supplementary Figure 1). The extent of these changes varied considerably, ranging from 1,033 differentially abundant proteins in KCNT1-Related Epilepsy and 910 in Fragile X Syndrome to fewer than 10 in DLG4-Related Synaptopathy and SLC6A1-Related Disorder (Figure 2B, Supplementary Table 1). Across disorders, proteins were more frequently increased than decreased (1,553 versus 931; 62.5%, binomial *p* = 5.4 x 10^-36^) (Figure 2C, Supplementary Table 2).

**Figure 2.**
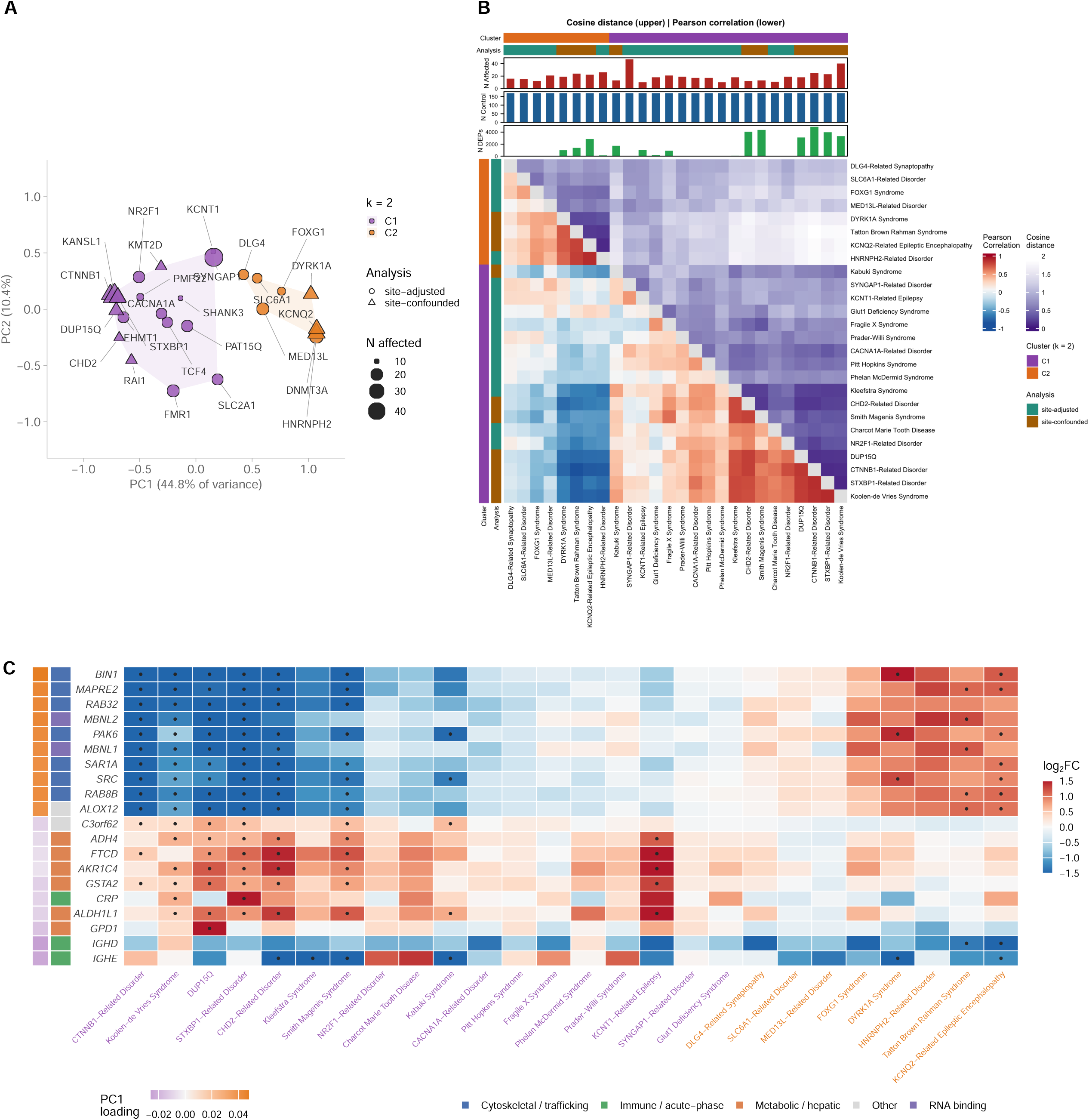
Differential plasma protein abundance across 26 rare genetic neurodevelopmental disorders. Differential abundance was assessed per disorder against all available unaffected controls (n = 170) using limma with empirical Bayes moderation, modeling log2(RFU) ∼ Group + Sex + Age. For the 16 disorders in which study site was identifiable, site was included as a random effect (duplicateCorrelation, block = Site); for the remaining 10, disorder and site are collinear and no site term was fitted (Methods). Proteins were called differentially abundant at Benjamini-Hochberg FDR < 0.10 with |log2 fold-change| > 0.2, corrected within disorder across 10,738 analytes. (A) Analysis paths, showing how the identifiability of study site determines which of the two analyses a disorder enters and what each supports. (B) Volcano plots for three site-adjusted disorders spanning the observed range of perturbation. Each point is one analyte; red and blue mark proteins passing the FDR and fold-change criteria in each direction, gray the remainder. The y axis shows nominal -log10(p) and the dashed horizontal line marks the p-value of the least significant protein passing FDR < 0.10, so every point above that line is FDR-significant; the axis is not -log10(FDR) because Benjamini-Hochberg produces extensive ties that collapse the vertical spread (Supplementary Methods). Vertical dashed lines mark the fold-change threshold. (C) Proteins differentially abundant in at least three site-adjusted disorders, with a functional annotation strip derived from GO enrichment of the same set. Rows are hierarchically clustered; the dot overlay marks disorder-protein pairs passing FDR < 0.10.

Recruitment site strongly influenced the apparent size of these proteomic signatures. The 10 disorders in which diagnosis and recruitment site were confounded showed substantially more differentially abundant proteins, consistent with site-associated variation rather than a greater biological effect. We therefore analyzed site-adjusted and site-confounded disorders separately throughout the study (see Method section). The overall pattern was consistent across different significance thresholds but was more strongly affected by how recruitment site was accounted for (Supplementary Figure 2A-D). These findings show that most RGNDs have detectable plasma proteomic signatures, but that the magnitude of these signatures can only be reliably interpreted when the recruitment site is accounted for.

Age also influenced RGND-associated proteomic signatures. Significant disorder-by-age interactions were detected in all 16 site-adjusted disorders, involving 2,602 proteins in total, with age-dependent effects also observed across all 10 site-confounded disorders (Supplementary Figure 2E, 2F). KCNT1-Related Epilepsy showed the largest number of age-dependent changes (1,461 proteins), despite participants spanning only 0-9 years of age, indicating substantial proteomic variation even within childhood. Pitt Hopkins Syndrome (268 proteins; ages 1-38 years) and FOXG1 syndrome (171 proteins; ages 2-20 years) also showed substantial age-dependent effects across broader age ranges. Together, these findings show that age modifies disease-associated proteomic signatures across RGNDs. Notably, when age was modeled directly, restricting the analysis to pediatric participants did not increase the number of differentially abundant proteins, as previously suggested (Supplementary Table 3).

Individual disorder signatures also revealed proteins with possible links to the underlying disease biology. Elevated CREG1 and CTSZ in Koolen-de Vries syndrome pointed to altered autophagy^5,6^, consistent with the known function of KANSl1^7^. Reduced STX1A in STXBP1-Related Disorder implicated presynaptic vesicle function^8^, while elevated TXNDC11 in Tatton-Brown-Rahman syndrome suggested endoplasmic reticulum stress^9^. Elevated NOGGIN in DYRK1A Syndrome pointed to altered BMP signaling^10^, and elevated PCDHB8 in SLC6A1-Related Disorder was consistent with altered synaptic organization^11^. In contrast, no disorder-specific candidate emerged for Fragile X Syndrome or Phelan-McDermid syndrome, where the cognate protein itself may provide a more informative marker. Although these candidates require validation in independent cohorts, they provide testable links between causal genetic changes and downstream disease biology.

Together, these results show that most RGNDs have detectable plasma proteomic signatures that vary across disorders and with age, while individual protein changes point to disease-relevant biological pathways.

### Genetically distinct disorders converge into two proteomic groups

Although each RGND showed its own plasma proteomic signature, we asked whether genetically distinct disorders converge on shared molecular patterns. Comparing proteomic changes across disorders revealed two groups, C1 comprising 18 disorders and C2 comprising 8 disorders (Figure 3A-C). The two-group structure was supported by multiple independent clustering approaches (Supplementary Figure 3A-C; Supplementary Table 4) and remained consistent across alternative analytical approaches (Supplementary Figure 3D). Removing each disorder individually reproduced the same grouping for 22 of 26 disorders, indicating that no single disorder drove the separation. Importantly, the grouping was independent of the site-adjusted and site-confounded analyses (Fisher’s exact p = 1.00), further supporting that the two-group structure reflects shared proteomic biology rather than the analysis design (Supplementary Figure 3E).

**Figure 3.**
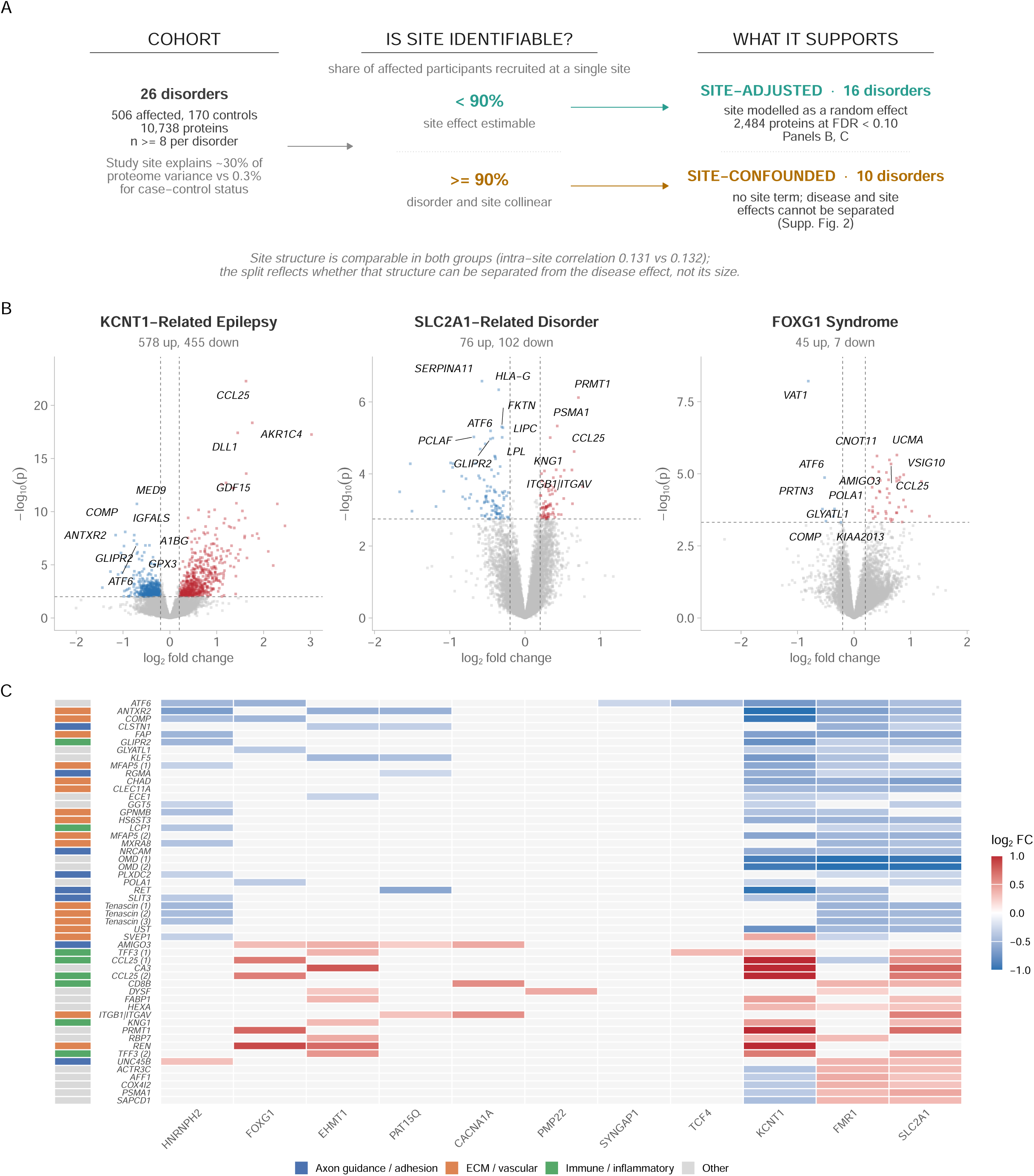
Cross-disorder proteomic similarity resolves two clusters along a single dominant axis. (A) Principal component analysis of per-disorder log2 fold-change profiles, unit-normalized so that position reflects the direction rather than the magnitude of each disorder’s proteomic change; on raw profiles PC1 correlates with the number of differentially abundant proteins (r = -0.65) and would therefore display effect size rather than similarity. Point size is the number of affected participants and shape denotes the analysis a disorder enters. (B) Loadings of the leading proteins on PC1, colored by pole. (C) Pairwise similarity between disorders: cosine distance above the diagonal and Pearson correlation below, both invariant to effect magnitude and in close agreement (Mantel R = 0.999). Annotation tracks give group membership, the analysis each disorder enters, and per-disorder sample and protein counts. (D) Proteins ranked by the difference in mean log2 fold-change between the two clusters, shown across all disorders ordered by PC1 score, with strips giving the signed PC1 loading and GO-derived functional class. Dots mark disorder-protein pairs passing FDR < 0.10.

Some proteomic changes were shared across both groups, suggesting common features of RGND biology. For example, ATF6, a regulator of the endoplasmic reticulum stress response^12^, and cartilage oligomeric matrix protein (COMP) were consistently reduced across both groups. These recurrent changes may reflect molecular features shared broadly across RGNDs rather than features that distinguish C1 from C2.

The two groups separated completely along the dominant proteomic axis, which accounted for 44.8% of the variation (Figure 3). Proteins contributing most strongly to this separation represented two broad biological classes. Cytoskeletal and vesicle-trafficking proteins, including BIN1, MAPRE2, RAB32, SRC and PAK6, characterized one end of the axis, whereas metabolic proteins, including ADH4, AKR1C4, FTCD, ALDH1L1 and GSTA2, characterized the other and were enriched for primary alcohol, folate and modified amino acid metabolism.

Notably, these proteins were not unique to either group. Rather, C1 and C2 differed primarily in the direction and magnitude of changes in shared proteins, suggesting that the two groups represent different positions along a common proteomic axis rather than completely distinct molecular pathways (Supplementary Table 5).

### Shared proteomic changes reveal epilepsy-associated and druggable proteins

Having identified shared proteomic patterns across disorders, we next asked which individual proteins recur across genetically distinct RGNDs. Among the 16 site-adjusted disorders, 40 proteins were differentially abundant in at least three disorders with at least 70% directional consistency. This recurrence was substantially greater than expected by chance: permutation analysis identified a median of only 6 recurrent proteins and a maximum of 14 across 1,000 permutations, compared with the 40 observed (6.6-fold enrichment, *p* < 0.001). Because site-confounded disorders showed substantially greater recurrence attributable to site-associated variation, subsequent cross-disorder analyses focused on the 40 proteins identified in the site-adjusted disorders (Supplementary Tables 6 and 7). These findings indicate that recurrent protein changes across RGNDs represent a shared proteomic signal rather than the expected overlap among individual disorder signatures.

We next asked whether this cross-disorder convergence extended to a shared clinical feature. Because epilepsy is a common comorbidity across RGNDs, we evaluated whether a shared plasma signature of epilepsy could be identified independent of genetic diagnosis. Across 13 site-adjusted disorders in which epilepsy status varied among affected participants, random-effects meta-analysis (REML) identified 32 epilepsy-associated proteins, of which 21 were increased and 11 decreased in participants with epilepsy (Figure 4A). The strongest associations included coproporphyrinogen oxidase (CPOX; FDR = 0.005), retinol binding protein 4 (RBP4; FDR = 0.006), GINS4, nidogen-1 (NID1) and carboxypeptidase M (CPM). Effects were highly consistent across disorders (median I² = 0; maximum = 31%), supporting a shared epilepsy-associated signal rather than an effect driven by a single diagnosis. Six of the 32 proteins are targets of approved drugs (Supplementary Table 8), indicating that components of this shared epilepsy signature are already pharmacologically accessible.

**Figure 4.**
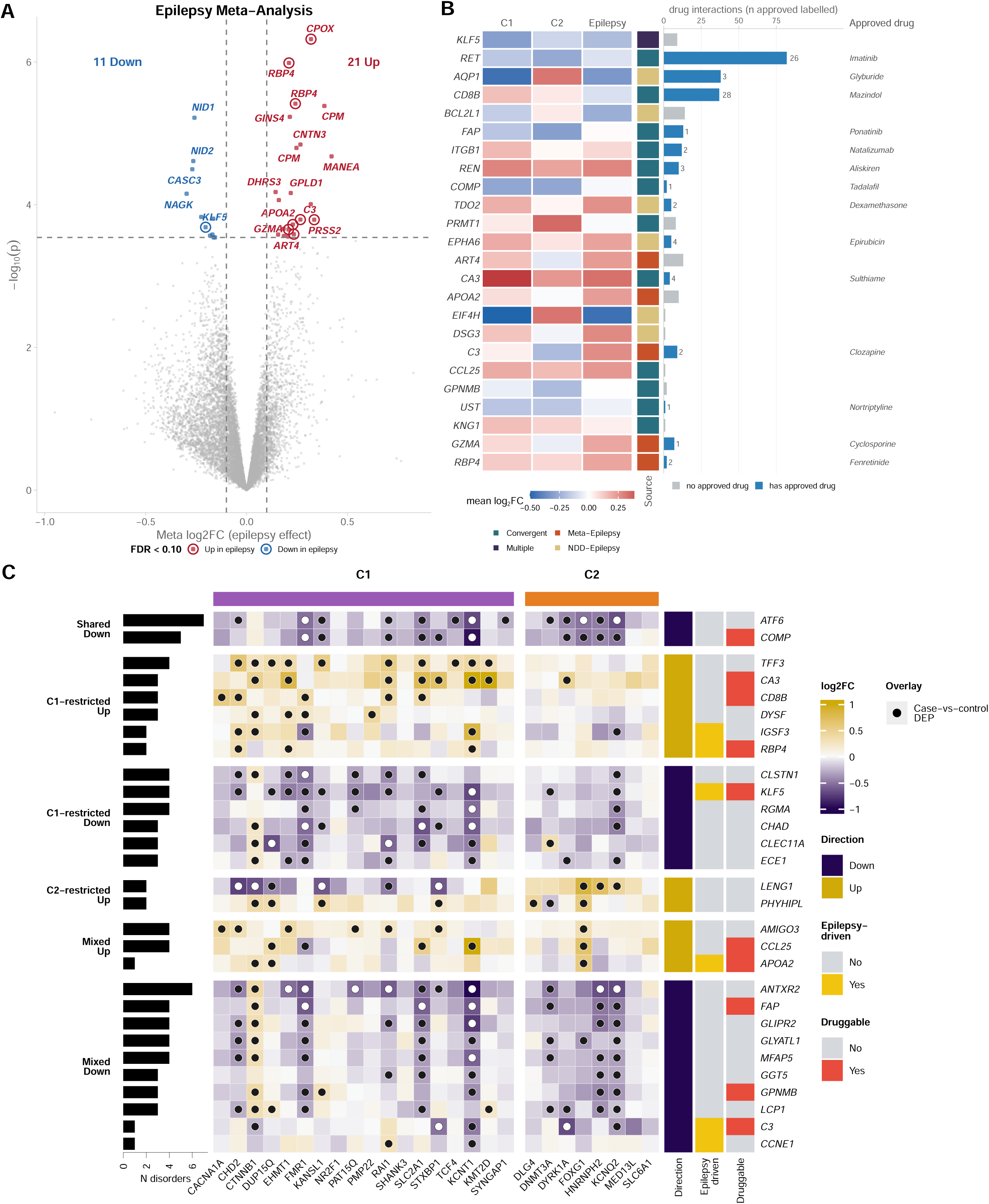
Epilepsy-associated plasma proteins and druggable targets across rare genetic neurodevelopmental disorders. (A) Volcano plot of epilepsy-associated proteomic changes from a random-effects meta-analysis (metafor, REML) of within-disorder epilepsy contrasts across the 13 site-adjusted disorders in which epilepsy status varies among affected participants. Standard errors were derived from limma moderated t-statistics as |log2FC/t|, and proteins were retained where at least five disorders contributed an estimate; 10,698 proteins met this criterion. Thirty-two proteins reached Benjamini-Hochberg FDR < 0.10 with |meta log2FC| > 0.10, of which 21 were higher and 11 lower in participants with epilepsy. Circled points are targets of an approved drug. Heterogeneity was negligible across the significant set (median I2 = 0, maximum 31%). (B) Druggable target integration for 24 nominated proteins. Tiles give the mean log2 fold-change within each cluster (C1, C2) and the pooled epilepsy effect; the adjacent strip gives the evidence source that nominated each target. Bars show the number of drug interactions recorded in DGIdb, colored by whether at least one approved drug exists, with the count of approved drugs labeled and a representative agent named at right. Endogenous compounds, nutrients and laboratory reagents returned by DGIdb with an approved flag were excluded from the approved-drug count. (C) Biomarker summary classification of 29 prioritized proteins across the 26 disorders, grouped by whether they recur in both clusters in the same direction (Shared, n = 2), in one cluster only with no contribution from the other (C1-restricted, n = 12; C2-restricted, n = 2), or in both clusters but below the recurrence threshold in one (Mixed, n = 13). Within each block proteins are ordered by dominant direction. Right-hand strips give direction, epilepsy association and druggability. Dots mark disorder-protein pairs passing FDR < 0.10

Notably, most epilepsy-associated proteins did not reach significance within individual disorders. Across the 32 proteins, only 21 of 840 disorder-protein comparisons reached significance within a single disorder, and 16 of the 30 proteins with the strongest pooled evidence were not significant in any individual disorder. Thus, combining evidence across genetically distinct RGNDs revealed a shared epilepsy-associated signature that would largely be missed when disorders are analyzed individually.

Finally, we integrated cross-disorder recurrence, proteomic group membership, epilepsy association and druggability to identify potentially druggable proteins (Figure 4C). Of 29 prioritized proteins, two recurred in both C1 and C2 in the same direction, 12 were specific to C1, two to C2 and 13 showed mixed patterns. Twenty-four of the 29 proteins have at least one recorded drug interaction, including targets of approved agents such as everolimus (AKT1), citalopram (GSK3B) and lithium citrate (GSK3A) (Supplementary Table 9). Together, these findings identify recurrent and epilepsy-associated proteins shared across genetically distinct RGNDs, many of which are already pharmacologically druggable.

### Rare neurodevelopmental disorders share proteomic features with adult neurodegeneration

We next asked whether the shared proteomic changes observed across RGNDs extend to adult neurodegenerative diseases. Comparing each disorder with reference plasma signatures of Alzheimer disease, Parkinson disease, frontotemporal dementia and amyotrophic lateral sclerosis revealed widespread overlap (Figure 5A). Seventy-one of 104 disorder-disease comparisons were significantly enriched, and 25 of 26 RGNDs showed enrichment for at least one neurodegenerative disease signature (Figure 5B). Frontotemporal dementia showed the broadest overlap, with enrichment in 21 of 26 disorders, followed by Parkinson disease (19), amyotrophic lateral sclerosis (17) and Alzheimer disease (14). Notably, the overlap was more often inverse than concordant: 46 significant comparisons showed negative enrichment compared with 25 showing positive enrichment. Six RGNDs - CTNNB1-, DNMT3A-, DYRK1A-, HNRNPH2-, KCNQ2- and KMT2D-related disorders - were enriched for signatures of all four neurodegenerative diseases. Similar enrichment patterns were observed at the proteomic group level (Supplementary Figure 4A; enrichment scores for all 104 comparisons, Supplementary Table 10). Proteins altered across RGNDs also showed substantial overlap with proteins altered in the reference neurodegenerative signatures and those supported by Mendelian randomization (Supplementary Figure 4B). Thus, RGNDs and adult neurodegenerative diseases share broad proteomic features, although these changes more often occur in opposing directions.

**Figure 5.**
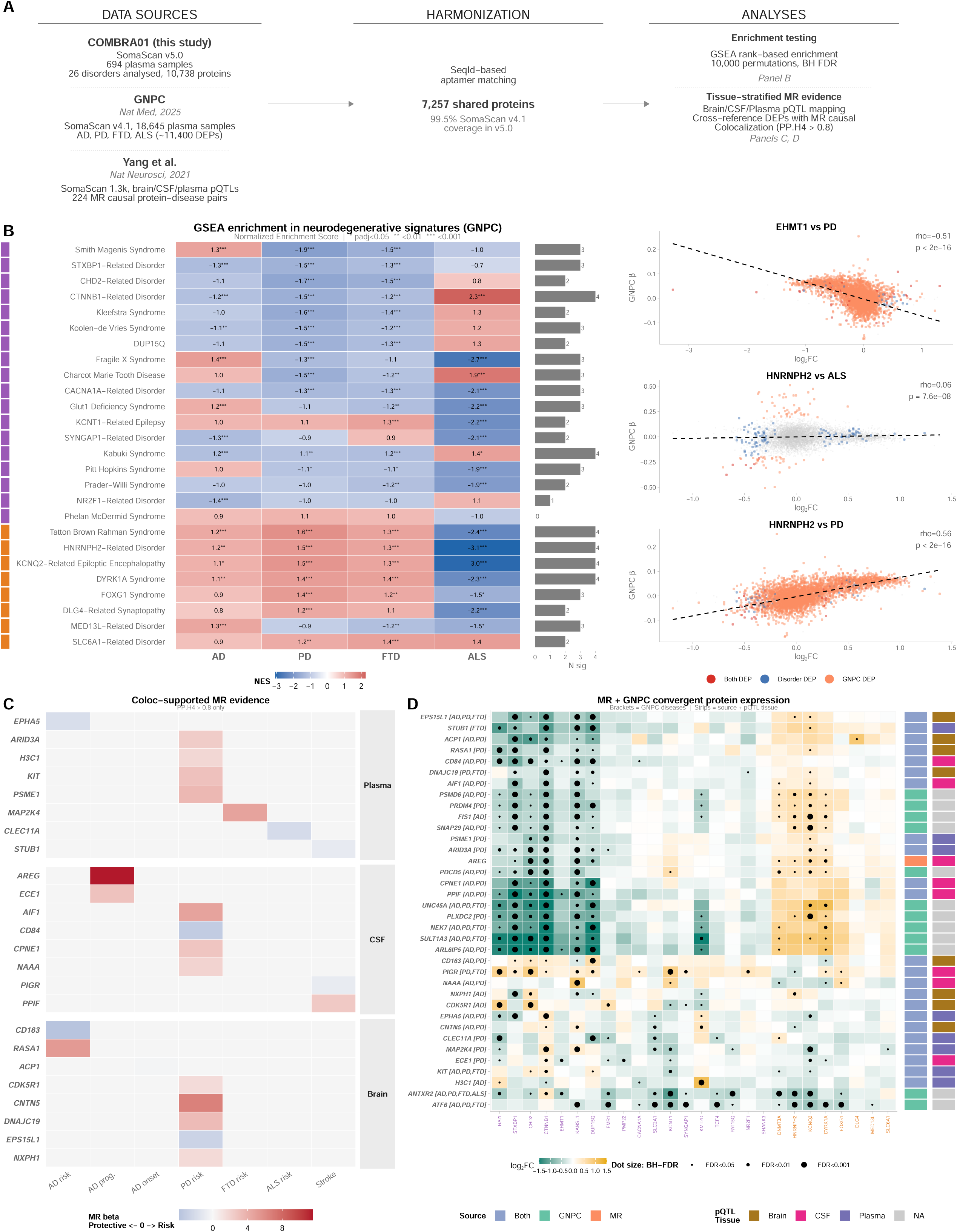
Cross-study convergence of rare neurodevelopmental disorder proteomes with neurodegenerative disease biology. (A) Study design and analytical workflow. Schematic illustrating the three data sources (COMBRA01, GNPC, Yang et al.), SeqId-based harmonization, and downstream analyses: GSEA enrichment testing (Panel B), tissue-stratified MR mapping (Panel C), and convergent protein integration (Panel D). (B) Gene set enrichment of neurodegenerative signatures across disorders. Left: normalized enrichment scores for each of 26 disorders against four reference signatures (104 tests), with 71 significant at Benjamini-Hochberg FDR < 0.05; asterisks denote adjusted significance and the strip at left gives cluster. Right: representative scatter plots of disorder log2 fold-change against reference effect size for three site-adjusted disorders, with Spearman coefficients inset. (C) Tissue-stratified Mendelian Randomization evidence. Tile heatmap of colocalization-supported (PP.H4 > 0.8) MR proteins from Yang et al. that are DEPs in ≥2 RGNDs, faceted by pQTL tissue of origin (Plasma, CSF, Brain). Within each tissue panel, proteins (rows) are shown across 27 disorders (columns); fill color encodes log2 fold-change direction (blue = downregulated, red = upregulated). Black dots indicate nominal significance (p < 0.05). Proteins are ranked by number of disorders in which they are differentially expressed. (D) Convergent proteins linking rare neurodevelopmental disorders to adult neurodegeneration. Heatmap of the 36 proteins supported by proteomic enrichment against a reference neurodegenerative signature, by Mendelian randomization evidence, or by both (23 by both, 12 by enrichment alone and 1 by genetic evidence alone), shown across the 26 disorders. Dot size denotes significance at FDR < 0.10; right-hand strips give evidence source and the tissue in which the causal association was detected.

We then examined whether proteins shared between RGNDs and adult neurodegenerative diseases also had independent genetic evidence linking them to neurodegeneration. Among 224 protein-disease pairs with colocalization support (posterior probability of a shared causal variant > 0.8), 79 unique proteins were also differentially abundant in at least one RGND (Figure 5C). These included proteins supported by plasma (34), cerebrospinal fluid (24) and brain tissue (24) pQTL evidence. Amphiregulin (AREG) was altered across nine RGNDs, while polymeric immunoglobulin receptor (PIGR) and ARID3A were each altered across eight. Importantly, this genetic evidence supports a potential causal role for these proteins in adult neurodegenerative diseases but does not establish causality in the RGNDs studied here (Supplementary Table 11).

Finally, integrating proteomic enrichment with Mendelian randomization evidence^13^ identified 36 proteins linking RGNDs with adult neurodegeneration (Figure 5D). Twenty-three were supported by both proteomic enrichment and genetic evidence, 12 by enrichment alone and one by genetic evidence alone. Only two proteins, endothelin converting enzyme 1 (ECE1) and C-type lectin domain containing 11A (CLEC11A), overlapped with the recurrent cross-disorder proteins identified in Figure 4. This limited overlap suggests that proteins recurrently altered across RGNDs are largely distinct from those linking RGNDs to adult neurodegeneration (Supplementary Table 12).

### A plasma proteomic signature tracks adaptive communication severity across disorders

We next examined whether plasma proteomic variation tracks how severely individuals are affected across RGNDs. General neurodevelopmental skills were assessed using the composite domain standard scores for the Vineland Adaptive Behavior Scales, Third Edition, Parent/Caregiver Form (Vineland-3; VABS-III)^14^, (Figure 6; Table 1). Developmental skills varied widely across disorders. Mean composite scores were at least two standard deviations below the normative mean for all disorders except Kabuki Syndrome, Prader-Willi Syndrome, Smith-Magenis Syndrome, with several disorders showing substantially greater impairment. Receptive and Expressive Communication scores were lowest in KCNT1-Related Epilepsy and FOXG1 Syndrome, identifying these as the two disorders with the greatest communication impairment in the cohort.

**Figure 6.**
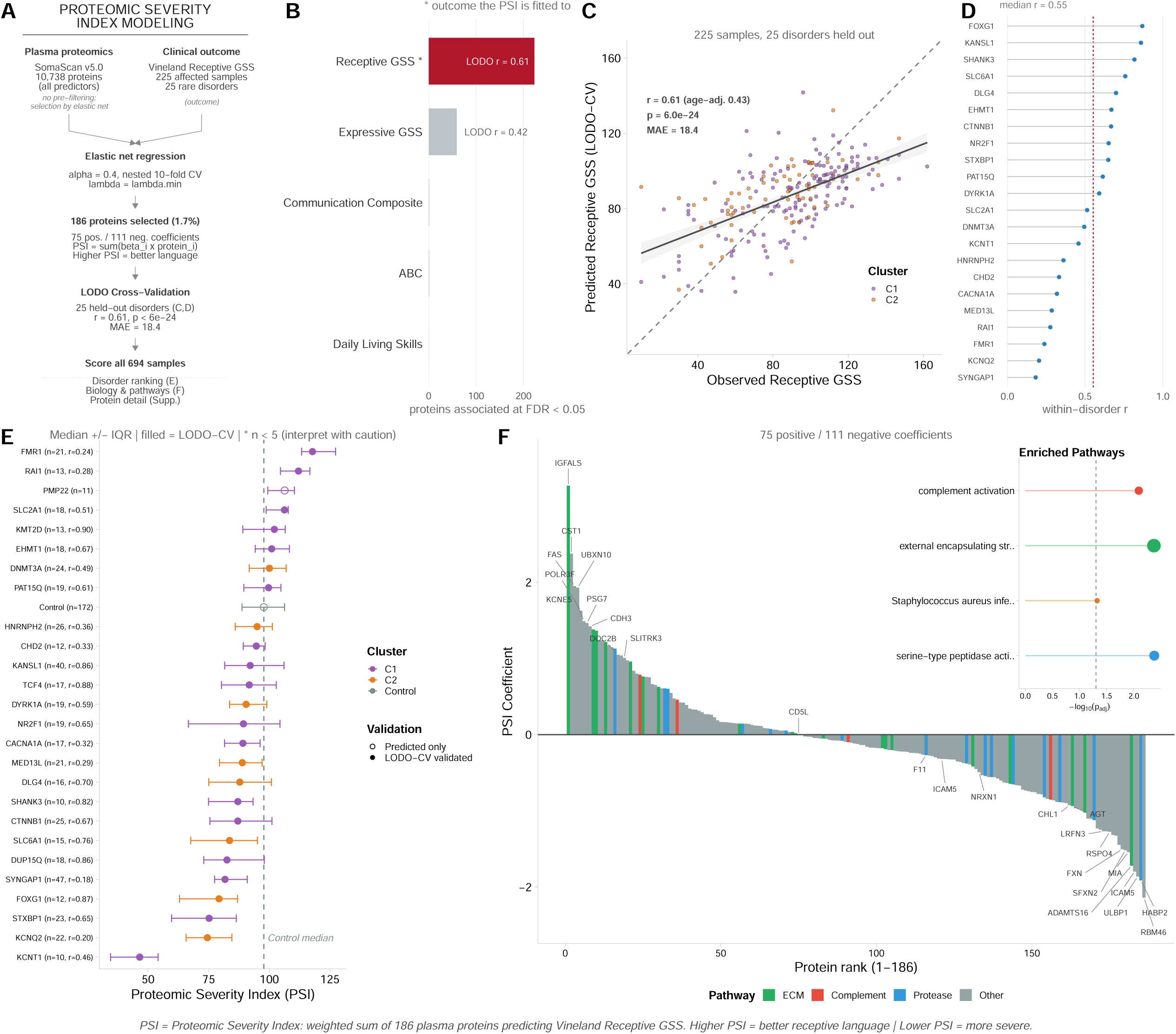
Proteomic Severity Index predicts clinical severity across rare genetic neurodevelopmental disorders. (A) Model development. Elastic net regression (alpha = 0.4, nested 10-fold cross-validation, lambda.min) was trained on all 10,738 SomaScan analytes without pre-filtering to predict the Vineland-3 Receptive Communication Growth Scale Score, a measure of adaptive communication behavior, in 225 affected participants across 25 disorders. The model retained 186 proteins (1.7%), 75 with positive and 111 with negative coefficients. The Proteomic Severity Index (PSI) is the weighted sum of these proteins; higher PSI indicates better receptive communication and lower PSI greater severity. (B) Outcome specificity. The same pipeline applied to each Vineland domain in turn yields 223 severity-associated proteins at FDR < 0.05 for Receptive Communication GSS, against 59 for Expressive Communication and at most one for the Adaptive Behavior Composite, Communication Composite and Daily Living Skills, indicating that the association is specific to the fitted outcome rather than a general property of a wide predictor matrix. (C) Leave-one-disorder-out cross-validation: each disorder was held out in turn, the model refitted on the remainder, and the held-out participants predicted. Observed against predicted Receptive GSS gives r = 0.61 (p = 6.0 x 10-24, mean absolute error 18.4) across 225 samples and 25 held-out disorders; color denotes cluster. Because the Vineland GSS is a growth scale score that rises with development, the age-adjusted correlation is also given (r = 0.43); the PSI adds information beyond age alone (incremental R2 = 0.130, nested F test p = 1.6 x 10-11). (C) Leave-one-disorder-out cross-validated prediction of Receptive GSS, each point one participant in a held-out disorder (r = 0.61, age-adjusted 0.43; mean absolute error 18.4). (D) Within-disorder correlation between observed and predicted Receptive GSS for the 22 disorders with at least five participants; the median is r = 0.55 against a between-disorder correlation of 0.81, indicating that the PSI tracks severity among individuals who share a diagnosis rather than only ranking disorders against one another. (E) Forest plot of median PSI with interquartile range across disorders and controls; filled points denote disorders validated in leave-one-disorder-out cross-validation and asterisks disorders with fewer than five participants, which should be interpreted with caution. (F) Coefficient waterfall of all 186 PSI proteins ordered from most positive to most negative, colored by pathway class, with the leading proteins labeled. Inset: Gene Ontology biological process terms enriched among the selected proteins. The heatmap of the top 40 PSI proteins by absolute coefficient is shown in Supplementary Figure 5E. Note that the color scale in that panel is the PSI regression coefficient per standard deviation of Receptive GSS, not a case-control fold change.

At the individual protein level, Receptive Communication showed substantially more severity-associated proteins than Expressive Communication across the proteome (223 versus 59 FDR-significant proteins; Supplementary Figure 5A,B), motivating its use for the cross-disorder proteomic index. Among proteins overlapping with adult neurodegeneration, 48 of the 50 proteins most strongly associated with severity reached FDR significance across disorders, including UCMA in STXBP1-Related Disorder, thrombin (F2) in Fragile X syndrome and AKR1C3 in DLG4-Related Synaptopathy.

We next asked whether the plasma proteome could capture communication ability across genetically distinct RGNDs. A proteomic index based on Receptive Communication Growth Scale Score retained 186 proteins (Figure 6A). When each disorder was held out in turn, the index correlated with observed Receptive Communication scores across 225 participants and 25 held-out disorders (r = 0.61, p = 6.0 × 10^-24^; mean absolute error = 18.4; Figure 6C). Importantly, this relationship was not explained by age: the association remained after age adjustment (r = 0.43), and the proteomic index explained additional variation beyond age alone (incremental R² = 0.135, p = 1.6 × 10^-11^). The index also tracked differences in severity among individuals with the same diagnosis, with a median within-disorder correlation of 0.55 across 22 disorders with at least five participants, compared with a between-disorder correlation of 0.81 (Figure 6D; per-disorder performance, Supplementary Figure 5C). Consistent with the individual-protein analyses, Receptive Communication showed a stronger proteomic association than the other Vineland-3 domains (Figure 6B; Supplementary Table 13). Together, these results show that plasma protein levels track communication severity beyond differences attributable to age or diagnosis.

The 186 proteins contributing to the index also pointed to biologically relevant pathways. They were enriched for extracellular matrix organization, complement activation and serine-type peptidase activity (Figure 6F, inset), consistent with established roles for extracellular matrix remodeling in synaptic organization^15^ and complement-mediated developmental synaptic pruning^16^. The strongest positive predictor was IGFALS, a component of the IGF complex involved in somatic and brain growth^17^. Other selected proteins included ICAM5, FXN, LRFN3, DOC2B and SLITRK3, which have established roles in neuronal and synaptic biology^18–22^. Five proteins had independent Mendelian randomization evidence for effects on brain phenotypes, and thirteen have established clinical laboratory assays (Figure 6F; proteins with the largest coefficients and their per-disorder associations, Supplementary Figure 5E). At the disorder level, median index values reflected differences in severity, with more severely affected disorders, including KCNT1-Related Epilepsy and KCNQ2-Related Epileptic Encephalopathy, separated from disorders with scores closer to the control median (Figure 6E).

Finally, we asked whether proteins associated with severity overlapped with those identified in our earlier cross-disorder analyses. The 186 index proteins showed modest overlap with proteins altered in neurodegenerative disease (125 of 186; 67%; hypergeometric p = 0.05), although this enrichment was not significant after correction across the four frameworks tested. In contrast, none overlapped with the recurrent cross-disorder proteins from Figure 4, the epilepsy-associated protein set or the convergent RGND-neurodegeneration proteins from Figure 5D (Supplementary Figure 5D; Supplementary Table 14). Thus, proteins that track communication severity represent a largely distinct proteomic signal from those that distinguish or recur across disorders.

Together, these findings show that plasma protein levels track differences in global developmental skills across RGNDs beyond the effects of age or diagnosis, while highlighting roles for IGF signaling, extracellular matrix remodeling, complement activation and synaptic biology. Because the index was developed and evaluated within a single cross-sectional cohort, it reflects current developmental skills rather than prospective prediction and will require validation in an independent cohort.

## Discussion

Genetically distinct rare neurodevelopmental disorders converge on a shared plasma proteome that overlaps the proteomes of adult neurodegenerative disease, and that carries graded information about how severely an individual is affected. Together these findings reframe how RGNDs are conceptualized and how they might be targeted therapeutically. Genetically distinct RGNDs converge onto an overlapping set of shared proteomic programs, organized along a dominant axis that separates two clusters, suggesting shared downstream biology rather than distinct gene-specific etiologies. The shared molecular overlap highlights the feasibility of aggregating RGNDs biospecimens that lack statistical power alone, but when combined unveil signatures, nominate biomarkers for consideration for clinical trial design, and can inform on future therapeutic efficacy studies. These observations collectively argue for a clinical trial paradigm in which: (i) entry is still based on precise genetic diagnosis; but (ii) outcome panels are pathway centric and shared across multiple proteins; and (iii) severity indices like PSI provide a common language to compare efficacy between conditions. Such designs could substantially increase statistical power and feasibility in ultra rare disorders by allowing data pooling across mechanistically related syndromes while still respecting genetic specificity for inclusion criteria and safety considerations.

Aggregating across disorders identified proteins recurring far more widely than any single-disorder study could show. The clearest is ATF6, reduced in seven of the 16 site-adjusted disorders, implicating the endoplasmic reticulum stress response as a shared axis. Others recur across the cohort but reach the convergence threshold only when the site-confounded disorders are included (Supplementary Figure 6A), among them GDF15, CCL11 and HSD11B1, each significant in one or two site-adjusted disorders; these are reported as exploratory rather than as established pan-RGND markers, and together they point towards glucocorticoid and neuroinflammatory pathways. Their support is nonetheless directional rather than absent: across all 16 site-adjusted disorders, and irrespective of whether any individual comparison reaches significance, GDF15, CCL11 and HSD11B1 each shift upward in 14, and ATF6 and COMP downward in 15, of the 16 (Supplementary Figure 6B, 6C). A skew of this size is not on its own unusual, since 688 of 10,738 analytes reach at least 14 of 16, so it is offered as a shared trend consistent with the disorder-level results rather than as independent evidence of convergence. Elevated GDF15 across RGNDs is particularly intriguing as it has been proposed as a biomarker for diabetes, mitochondrial disorders, cancer and several neurological diseases attributed to neurodegeneration and neuronal injury^23–27^. More recently, elevated plasma GDF15 was found to be predictive of brain aging - using multimodal imaging^28^. Plasma GDF15 was found to be specifically linked to dementia risk, but perhaps more relevant to RGNDs, elevated GDF15 was correlated with poorer executive function skills and impaired motor function^29^. The molecular functions of GDF15 are multifaceted and include cellular senescence as a primary role - and ultimately a regulator of neuroinflammation, neuronal injury and apoptosis^30^. This pattern is consistent with convergent inflammatory and matrisome dysregulation described across neurodegenerative diseases^31^, where distinct primary etiologies nonetheless converge on overlapping immune and synaptic pathways. Additionally, GDF15 and related senescence markers may provide initial insights into the overlap between RGNDs and AD pathology compared to other neurodegenerative disorders.

Age exerts disorder specific effects on the systemic proteome, with some syndromes showing stronger perturbation in childhood while others exhibit abnormalities into adulthood, re-shaping biomarker consideration for pediatric versus adult trial design. In neurotypical aging, more than 200 plasma proteins correlate significantly with age, many related to immune function and inflammatory processes^28^. Age further modulated proteomic signatures in a disorder-specific manner. Fitting a group by age interaction within a single model per disorder, rather than comparing counts between separately fitted pediatric and all-ages models, showed FDR-significant interactions in every disorder analyzed, most extensively in KCNT1-Related Epilepsy, TCF4-Related Disorder and FOXG1 syndrome. These findings align with emerging large-scale neurodegeneration plasma proteomics studies, which shows that disease stage and age strongly shape circulating proteomic profiles and pathway enrichment patterns^28^. More importantly, these findings suggest that plasma biomarkers from pediatric studies may differ from adult studies; this has relevance to upcoming RGND clinical trial design. In addition to age, variant type heterogeneity as well as pathogenic gene expression throughout development likely contribute to DEP variability across age.

If early-life monogenic disorders and late-onset neurodegeneration engage overlapping biology, that overlap should be visible in the plasma proteome. It is: RGND profiles overlapped extensively with signatures of adult neurodegenerative disease derived from large, independent cohorts. Most RGNDs showed significant enrichment with at least one neurodegenerative proteomic signature and many showed enrichment across several, with both concordant and inverse relationships. Integration with Mendelian randomization data further highlighted a set of causal neurodegeneration linked proteins that are also perturbed in RGNDs, particularly those observed in the CSF and point to immune response pathways as a shared substrate. Because these overlaps were identified by rank-based enrichment over the full ranked proteome rather than a thresholded list, they are not driven by differences in the number of differentially abundant proteins between disorders. That a childhood monogenic disorder and a late-onset degenerative disease can leave related signatures in plasma argues that the two are not as biologically separate as their clinical presentations suggest.

Mechanistically, this overlap places many RGNDs and common neurodegenerative conditions along a continuum of disrupted synaptic integrity, immune surveillance and cellular stress in the nervous system. Early life monogenic insults may set in motion proteomic programs that in later decades manifest as or intersect with pathways driving neurodegenerative conditions such as ALS, FTD, PD, or AD, consistent with emerging evidence that rare genetic disorders can illuminate generalizable mechanisms in brain aging and degeneration^32^. Conversely, compensatory or inverse enrichments raise the possibility that some RGND associated alterations might actually lower neurodegenerative risk trajectories, an idea that could be explored through longitudinal RGND natural history studies and registries. Taken together, this continuum supports a shift from single-gene to pathway-based clinical strategies, in which convergent molecular pathology shared with both other rare disorders and common neurodegenerative disease underpins cross-disorder biomarker frameworks and potentially shared treatments.

Clinically, these data carry three implications. First, they support the use of neurodegeneration oriented biomarker panels (e.g., those capturing complement, innate immune, and stress response proteins) in RGND natural history and interventional studies, enabling cross talk with larger neurodegeneration trials and biobanks. Second, they provide a rationale for cautious repurposing of neurodegenerative therapeutics—such as agents targeting complement, microglial activation, or integrated stress response—into RGND populations whose proteomes show concordant enrichment, provided age, safety, and developmental considerations are respected. Third, they raise the possibility that individuals with certain RGNDs may carry altered lifetime risk for neurodegenerative outcomes, either increased or decreased depending on the direction of proteomic overlap.

Epilepsy is the most common comorbidity across these disorders, and the meta-analysis identified proteins tracking seizure burden independent of genotype (CPOX, RBP4, GINS4, CPM), indicating that both genetic diagnosis and seizure burden sculpt the systemic proteome. Against that background, forty proteins met the cross-disorder convergence criterion within the site-adjusted analysis, recurring in at least three of 16 disorders with at least 70% directional consistency. Counting the same way within the site-confounded analysis returns 1,943 proteins, a 49-fold difference that reflects retained site-driven variation rather than greater biological convergence, which is why the two are reported separately and never pooled. Importantly, 24 of the 29 prioritized proteins have at least one recorded drug interaction, and a subset are also implicated in the epilepsy meta analysis, placing them at the intersection of disease mechanism, seizure burden, and druggability. This has promising clinical implications for RGNDs. First, these convergent markers provide immediate candidates for multi disease biomarker qualification: a small panel including ATF6, GDF15, CCL11, HSD11B1, and selected epilepsy associated proteins (e.g., CPOX, RBP4, CPM) could be deployed as a shared secondary endpoint across early phase trials spanning multiple RGNDs. Second, the Proteomic Severity Index (PSI) demonstrates that a relatively small subset of proteins, enriched for ECM and complement biology, can predict Vineland Receptive Communication Growth Scale Score, a measure of adaptive communication behavior, across 25 disorders, raising the possibility that transdiagnostic, proteomics based scores could be used to enrich for patients at particular severity ranges, to monitor progression, or to quantify the impact of interventions that aim to improve core neurodevelopmental function rather than a single gene’s expression. The potential to use the PSI to predict or evaluate developmental gains will be a critical tool in trial designs for RGNDs.

There are several important limitations to this study. The study is cross-sectional, precluding assessment of longitudinal biomarker trajectories or treatment response. Sample sizes varied substantially across disorders, from 10 (KCNT1-Related Epilepsy) to 47 (SYNGAP1-Related Disorder), limiting statistical power for the smallest cohorts; WWOX-Related Epileptic Encephalopathy (n = 6) fell below the eight-participant threshold and was excluded. Study site was a substantial source of proteomic variance and was modeled as a random effect wherever it was estimable, but in 10 of 26 disorders recruitment was so site-concentrated that disorder and site are collinear; those results are reported separately and descriptively and are never pooled with the site-adjusted analysis. Plasma proteins may also not fully recapitulate central nervous system pathology, and markers that track seizure proximity should not be interpreted as pure indices of central nervous system severity. This is measurable rather than hypothetical: among the 222 affected participants with epilepsy for whom a date of last seizure was recorded, the severity index rises with time since that seizure (age-adjusted partial correlation 0.29; Supplementary Figure 7A), and the association persists within disorders, though weakly (median 0.17 across the 12 disorders contributing at least eight such participants, Supplementary Figure 7B; permutation p = 0.012 against a within-disorder null centered at 0.19, Supplementary Figure 7C). Part of what the index captures is therefore proximity to recent seizure activity rather than developmental status alone. The metadata contain no medication, nutritional or inflammatory-comorbidity variables, so those adjustments could not be performed. The aptamer-based platform, while high-throughput, does not capture all circulating proteins and may exhibit non-specific binding to homologous proteins and target-specific binding variability. Vineland developmental data were available for only a subset of participants, restricting the severity model’s training set. Finally, leave-one-disorder-out cross-validation holds out whole disorders from the same cohort, sites and assay run, so it demonstrates generalization to an unseen disorder but is not external replication; independent replication in a separate cohort will be essential to validate the severity index and the cross-disorder grouping; per-disorder values are given in Supplementary Table 15.

Altogether, the convergent RGND clusters and their overlap with neurodegenerative proteomes reinforce the broader concept that clinically heterogeneous disorders can share core molecular pathways, and that therapies targeting those pathways can sometimes be redeployed across genetically distinct diseases. In RGNDs, this work points specifically toward shared targets in ER stress (ATF6 and downstream effectors), glucocorticoid and neurosteroid metabolism (HSD11B1, AKR1C3), complement and microglial signaling, where C3 and CXCL3 are reduced rather than elevated in the site-adjusted disorders in which they reach significance, and integrated stress response markers such as GDF15 that traverse developmental and degenerative conditions. The presence of existing drugs against many of these proteins, along with the ability of plasma proteomics to stratify patients and quantify both cross disorder biomarkers and individual severity, lays a foundation for rational, mechanism anchored repurposing and for multi arm trials that test shared interventions across genetically distinct but molecularly convergent rare diseases.

## Acknowledgements

We gratefully acknowledge the 27 rare disease patient foundations whose partnership made this work possible. These foundations include: The CACNA1A Foundation, Coalition to Cure CHD2, CTNNB1 Connect and Cure, Hereditary Neuropathy Foundation, DLG4 Research Fund, Hope for Harvey, Dup15Q Foundation, International DYRK1A Syndrome Association, FOXG1 Foundation, FRAXA, GLUT1 Deficiency Foundation, Yellow Brick Road Project, KCNQ2 Foundation, KCNT1 Epilepsy Foundation, Kabuki Syndrome Foundation, IDefine, Koolen-de Vries Syndrome Foundation, MED13L Foundation, NR2F1 Foundation, CureSHANK, Pitt Hopkins Syndrome Foundation, Foundation for Prader-Willi Research, SLC6A1 Connect, STXBP1 Foundation, Parents and Researchers Interested in Smith Magenis Syndrome, Cure SYNGAP1, Tatten Brown Rahman Syndrome Foundation, WWOX Foundation. Through their leadership, trust, and tireless engagement, these organizations mobilized their communities to participate in this study, enabling the scale and diversity of data essential to our findings. Their commitment to advancing research and improving outcomes for individuals and families affected by rare neurodevelopmental disorders continues to drive meaningful progress in the field. We are deeply appreciative of their collaboration and shared vision.

## Methods

### Recruitment

RGNDs subjects and unaffected siblings (see Table 1) were recruited by patient advocacy groups representing each of 27 disorders. Of these, 26 disorders had at least eight affected participants and were carried forward to differential abundance analysis; WWOX-Related Epileptic Encephalopathy (n = 6) fell below this threshold. Participants or caregivers provided consent via video teleconference with a COMBINEDBrain research coordinator Immediately after consenting, participants uploaded their genetic report to a HIPAA- and GDPR-compliant data collection platform, Across Healthcare’s Matrix. The genetic reports were curated by the research coordinators and principal investigator to confirm diagnoses. Caregivers were also asked to complete the Vineland Adaptive Behavior Scales - III Parent/Caregiver Form(VABS-III)^14^. The VABS-III is a standardized and norm-referenced measure of adaptive function and is commonly used as a proxy for developmental status. The Vineland-3 assesses three primary domains including Communication, Daily Living Skills and Socialization skills^14^. The Vineland-3 Adaptive Behavior Composite (ABC) and domain standard scores (all with m=100, SD=15) and Receptive and Expressive subdomain GSVs (Rasch-based person ability scores) were examined. At the time of sample collection, caregivers were asked about any recent hospitalizations and medication changes, when they had last eaten and any special dietary restrictions and when was their most recent seizure.

### Ethics

This study was conducted under ethical approval by NorthStar Review Board (#NB200141). Participants gave consent for de-identified samples and data to be used in research studies. All participants provided consent for sample donation into the COMBINEDBrain Biorepository. Further, for all the disorders included in this pilot project, a relevant patient advocacy foundation representing each disorder included in this pilot project provided approval for their communities’ patient samples to be included in this project.

### Blood Collection

Blood samples were collected at family and patient conferences, at home with a mobile phlebotomist, or in collaboration with a clinical or research lab. The research team confirmed participant consent and collected information related to diet, medications, time of last meal, and time of last seizure immediately prior to sample collection. The collection time was recorded for each blood sample prior to processing.

### Family Conferences

Research staff attended 17 scientific and family conferences throughout the United States between March and December 2023. Participants and caregivers diagnosed with any of the 27 represented disorders were eligible to attend any of the 17 conference collection sites. Conference-based blood collection for rare disorders feasible, bringing geographically dispersed families for each disorder to a single site.

### Home Visits

Mobile phlebotomists with expertise in pediatrics were contracted from Travalab, Speedy Sticks, and Modern Lab Services to complete residential blood collections. After consenting, a collection kit containing a 10ml EDTA tube, 15x 2ml cryovials, venipuncture supplies, a Data Collection Form to be completed by the phlebotomist was mailed to the participant prior to a mobile phlebotomy appointment. Mobile phlebotomists collected samples between the hours of 7am-4pm and processed blood according to the established protocol.

### Clinician Assisted

Some participants were actively enrolled in academic research studies which included biospecimen collections. In these instances, participants provided additional consent and were able to collect a blood sample for this study in addition to the primary research study. Samples were processed according to the processing protocol outlined below.

### Sample processing

A 10 mL EDTA tube (Fisher Scientific, 13-680-61) was collected via standard venipuncture and inverted 3-4 times to ensure EDTA distribution throughout whole blood. Samples were kept on ice prior to centrifugation (1500xg, 10 min, room temperature) to isolate plasma. Plasma was aliquoted into cryovials in 250 µl volumes and flash frozen on dry ice within 30-60 minutes of sample collection. Time from collection to centrifugation was recorded. Samples were stored at the COMBINEDBrain Biorepository at the Van Andel Institute (-80°C) until use. A plasma aliquot (250μl) was analyzed using SomaScan, a GAPmer-based technology, to measure 10,738 analytes, all of which were carried into the analyses reported here.

### Data Processing

The SomaScan® assay outputs raw intensity data as an .adat file, which contains relative fluorescence unit (RFU) values corresponding to the binding of SOMAmers to protein targets. Standard data processing and normalization procedures are performed to minimize technical variation and enable valid biological comparisons, as previously described9. The first step corrects for within-run hybridization variation. Hybridization control probes—exogenous sequences spotted on the array are used to adjust raw RFU values for systematic differences in hybridization efficiency across samples. To reduce systematic technical variability across samples within a run (e.g., pipetting differences, variation in assay handling), the median intensity across all SOMAmers in each sample is scaled to a reference distribution. This enforces global comparability of signal intensities. To compare data across assay plates or runs, normalization factors are calculated from reference calibrator samples (often pooled reference plasma or standardized controls) run on each plate. These factors are used to rescale sample values, correcting for batch effects and inter-plate variability. Quality control metrics are applied at the sample and SOMAmer level, including checks for signal range, coefficient of variation (CoV) in control replicates, detection outliers, and aptamer performance flags. Quality-control flags supplied by SomaLogic were reviewed, but no additional in-house exclusion was applied: all 694 samples and all 10,738 analytes in the delivered normalized dataset were retained for analysis. After normalization, log2 transformation of RFU values was applied to stabilize variance and approximate normal distributions prior to statistical modeling.

### Data Analysis

SomaScan proteomic data were analyzed using a standardized bioinformatics pipeline implemented in R. Raw relative fluorescence unit (RFU) values were log2-transformed. No additional in-house quality-control exclusion was applied beyond the normalization supplied by SomaLogic: all 694 samples and all 10,738 analytes in the delivered dataset were retained. Differential protein abundance was assessed per disorder using limma with empirical Bayes moderation^33^, modeling log2(RFU) as a function of case-control status, sex and scaled age (log2(RFU) ∼ Group + Sex + Age_scaled), with study site included as a random effect through limma duplicateCorrelation (block = Site)^33^ wherever site was estimable. Proteins were classified as differentially abundant at Benjamini-Hochberg adjusted FDR < 0.10 with |log2 fold-change| > 0.2, corrected within disorder across all 10,738 analytes. Cross-disorder similarity was assessed using cosine distance between disorder-level log2 fold-change profiles.

Study site and the two-tier analysis. Site accounted for a substantially larger share of proteome variance than case-control status and was therefore modeled as a random effect. This adjustment is not estimable for every disorder: recruitment was site-concentrated, and where a disorder’s affected participants were drawn overwhelmingly from a single site, disorder and site are collinear, so a site random effect absorbs the disease effect rather than adjusting for confounding. An identifiability criterion was pre-specified on the study design, before results were examined: a disorder’s site effect was treated as not identifiable when at least 90% of its affected participants were recruited at one site. This yields two analyses that are reported separately and never combined. The site-adjusted analysis comprises 16 disorders in which the site random effect is estimable. The site-confounded analysis comprises 10 disorders fitted without a site term; these estimate a combined disease-and-site effect that cannot be decomposed and are reported as descriptive only. Their differentially abundant protein counts are systematically larger precisely because they retain site-driven variation. Median consensus intra-site correlation is comparable in both clusters (0.131 and 0.132), so the distinction reflects identifiability rather than a difference in the amount of site structure. No combined differential-abundance total is reported, disorders are not ranked across the two analyses, and the two are not plotted on shared axes. All 26 disorders are retained in every other analysis, including cohort description, cross-disorder grouping, biomarker prioritization, neurodegeneration convergence and the severity index.

### Differential protein abundance analysis (Figure 1A-P, Figure 2A-C; Supplementary Figure 2A-F, Supplementary Figure 2)

For each of 26 disorders with at least eight affected participants, affected individuals were compared against all 170 available unaffected controls using limma with empirical Bayes moderation. Controls were not subsampled to a fixed case-control ratio; all controls contributed to every disorder comparison, with age and sex included as covariates (log2(RFU) ∼ Group + Sex + Age_scaled) and study site as a random effect where estimable. Proteins were classified as differentially abundant at Benjamini-Hochberg FDR < 0.10^34^ with |log2 fold-change| > 0.2. Age-dependence of the disease effect was assessed by fitting a group by age interaction within a single model per disorder, rather than by comparing the number of significant proteins between separately fitted all-ages and pediatric models, a comparison that cannot establish a difference. Cognate disease proteins (for example CTNNB1 for CTNNB1-Related Disorder) were identified using SeqId-to-gene mappings from the SomaScan manifest. Direction of change per disorder is given in Supplementary Table 2. Age modifies the disease effect in most disorders, with each protein assessed by an age x interaction model provided in Supplementary Table 3.

### Cross-disorder proteomic similarity and clustering (Figure 3A-C; Supplementary Figure 3A-E)

Disorder-level log2 fold-change profiles were computed across all 10,738 proteins for each of 26 disorders, generating a 26 by 10,738 matrix. Pairwise similarity was assessed using cosine distance, which depends on the direction rather than the magnitude of a disorder’s proteomic change; Euclidean distance between these profiles correlates with the number of differentially abundant proteins per disorder (r = 0.92) and therefore measures effect size rather than similarity. Hierarchical clustering (Ward.D2 linkage on cosine distance) defined the reported partition. The number of groups was evaluated using the gap statistic^15^, average silhouette width^35^ and consensus clustering over 1,000 bootstrap resamples^36^; k = 2 was selected, with k = 3 reported in the supplement. Profiles were embedded for display by principal component analysis on unit-normalized profiles rather than by uniform manifold approximation and projection, which is unstable at this number of points and whose layout depends on parameter choices. Robustness was assessed in three ways: agreement of the partition across four distance metrics and three linkages, leave-one-disorder-out clustering quantified by the adjusted Rand index^37^, and a Fisher exact test of independence between group membership and the site-adjusted or site-confounded analysis.

### Shared DEP identification (Figure 3D)

Proteins recurring across disorders were identified as those differentially abundant in at least three disorders with at least 70% directional consistency. Recurrence was tested against a permutation null in which analyte identities were shuffled while each disorder’s number of differentially abundant proteins was held fixed (1,000 permutations, seed 42), so the comparison isolates overlap between disorders from per-disorder burden. Representative proteins were ranked by the number of disorders showing significance and by mean absolute fold-change. Expression heatmaps display log2 fold-change capped at +/-1.5, with an overlay marking disorder-protein pairs passing FDR < 0.10.

### Epilepsy meta-analysis and druggable target identification (Figure 4A-C)

Convergent biomarkers and epilepsy meta-analysis hits were queried against the Drug-Gene Interaction Database (DGIdb)^67^ to identify proteins with existing pharmacological agents. Interaction records were retrieved per gene and summarized as the number of distinct drug interactions and the number of approved drugs. Endogenous compounds, nutrients and laboratory reagents returned by DGIdb with an approved flag (for example cholesterol, hydrogen peroxide, dimethyl sulfoxide and ascorbic acid) were excluded from the approved-drug count, since they do not represent therapeutic agents.

### Cross-study validation with neurodegenerative proteomics (Figure 5A-D; Supplementary Figure 4A, 4B)

Proteomic profiles from 26 rare neurodevelopmental disorders were cross-referenced with two external datasets: (1) the Global Neurodegeneration Proteomics Consortium (GNPC; Nat Med, 2025), which profiled approximately 18,645 plasma samples from individuals with Alzheimer’s disease (AD), Parkinson’s disease (PD), frontotemporal dementia (FTD), and amyotrophic lateral sclerosis (ALS) using SomaScan v4.1 (∼11,400 DEPs); and (2) Yang et al68, which identified 224 Mendelian Randomization (MR) causal protein–disease pairs from SomaScan 1.3k brain, CSF, and plasma pQTL data. Platform harmonization was performed at the SeqId level, yielding 7,257 shared proteins (99.5% coverage of v4.1 analytes in v5.0). GSEA enrichment was performed using the fgsea package76 with rank-ordered log2 fold-change profiles as input and GNPC disease DEP sets as gene sets (10,000 permutations; Benjamini–Hochberg FDR correction). Complementary Fisher’s exact tests assessed overlap enrichment, with significance confirmed by empirical permutation testing (10,000 permutations). Cluster-level enrichment was computed using Fisher’s combined p-value method (–2∑log(p) ∼ χ²). MR-supported proteins were filtered as those with colocalization support (PP.H4 > 0.843) and present as DEPs in ≥2 rare disorders. Effect size concordance was assessed by Spearman correlation of log2 fold-changes between RGND and GNPC DEPs. Inflammation sensitivity was evaluated by repeating Fisher tests after excluding 962 inflammation-associated genes. Epilepsy confounding was assessed by Spearman correlation between epilepsy prevalence and enrichment strength across disorders.

Convergent proteins from MR and GNPC evidence (Figure 5D) were subjected to GO Biological Process enrichment analysis using clusterProfiler::enrichGO^38^ with Benjamini–Hochberg FDR correction (q < 0.05). The 36 convergent proteins were defined as those supported by MR causal evidence (coloc-supported, PP.H4 > 0.8, DEPs in ≥2 RGNDs), GNPC enrichment (leading edge proteins from significant GSEA results), or both.

### Proteomic Severity Index (Figure 6A-F; Supplementary Figure 5A-D)

An elastic net regression model^39^ was trained to predict Vineland Adaptive Behavior Scales, Third Edition (Vineland-3) Receptive Communication Growth Scale Score (GSS) from plasma protein levels. Receptive GSS was selected as the outcome based on systematic comparison of five Vineland scores: the number of FDR-significant severity-associated proteins was quantified per score (global limma analysis, BH-corrected), and LODO-CV elastic net prediction accuracy was compared between Receptive and Expressive GSS (the two growth scale scores with sufficient dynamic range). The model used all 10,738 SomaScan v5.0 proteins as candidate predictors, enabling post hoc enrichment testing of biomarker frameworks. Optimal regularization was determined by grid search over 10 alpha values (0.1–1.0 in 0.1 increments) using nested 10-fold cross-validation; alpha = 0.4 minimized mean squared error. Lambda was selected at lambda.min via 10-fold CV (glmnet). The final model retained 186 proteins with non-zero coefficients (75 positive, 111 negative). Model performance was assessed using leave-one-disorder-out (LODO) cross-validation, in which each of the 26 evaluable disorders (≥3 samples) was sequentially held out and the model retrained on the remaining disorders; predicted values for held-out samples were compared to observed Receptive GSS. Performance metrics included Pearson correlation (r), mean absolute error (MAE), and significance of the observed–predicted correlation (cor.test). The Proteomic Severity Index (PSI) was defined as the model-predicted Receptive GSS for each individual, with lower PSI indicating greater clinical severity. Controls (n = 172) were never included in model training and served as zero-shot external validation. Hypergeometric enrichment tests assessed whether PSI proteins were over-represented among biomarker frameworks from Figures 4–5 (convergent biomarkers, epilepsy meta-analysis, Yang MR causal proteins, triple intersection, Fig 5D convergent, GNPC DEPs), using the full SomaScan proteome (N = 10,738) as universe; p-values were corrected using the Benjamini-Hochberg method across the six tests. Gene Ontology enrichment of PSI proteins was performed using clusterProfiler enrichGO (BP, MF, CC ontologies) and enrichKEGG, with the SomaScan proteome as background universe and BH-adjusted p < 0.05 as the significance threshold.

Severity-associated proteins were ranked by a combined score of global p-value and directional consistency (the proportion of disorders showing the same direction of effect). The top 50 GNPC-overlapping severity proteins were displayed in expression heatmaps with log2FC values capped at ±1.5 and significance annotations. Cross-framework enrichment of biomarker sets (Figure 4 convergent biomarkers, epilepsy meta-analysis, Yang MR causal proteins, GNPC DEPs, triple intersection) among the 186 PSI proteins was assessed using hypergeometric tests with the full SomaScan proteome (N = 10,738) as the background universe. The cross-framework evidence matrix (Figure 6E) displays PSI protein overlap with each biomarker framework for the top 15 proteins by absolute coefficient magnitude plus all proteins appearing in ≥2 frameworks.

## Software and visualization

All analyses were performed in R 4.4.1 (2024-06-14). Key packages: limma (differential expression and site random effects), metafor (random-effects meta-analysis), fgsea (rank-based gene set enrichment), glmnet (elastic net), clusterProfiler and org.Hs.eg.db (GO enrichment), ConsensusClusterPlus and mclust (clustering robustness), ComplexHeatmap and circlize (heatmaps), ggplot2, cowplot, and patchwork (figure assembly), and ggrepel (label placement). Drug-gene interactions were queried from DGIdb.

## Data availability

Individual-level proteomic and phenotypic data are available under controlled access through COMBINEDBrain. Open deposit is not possible: 427 of 522 affected participants (82%) are uniquely identified by the combination of disorder, age, sex and recruitment site, several disorders contribute a single participant, and each record links to a complete 10,738-analyte plasma proteome, so the dataset cannot be de-identified to a standard that would permit unrestricted release. Participants consented to research use of de-identified samples and data, with the advocacy foundation representing each disorder approving inclusion; that consent does not extend to open publication of individual-level profiles. Requests for access should be directed to COMBINEDBrain (combinedbrain.org) and will be considered under the terms of the original consent and the governing data use agreement. Summary-level results supporting every figure, including per-disorder differential abundance statistics, cluster assignments, enrichment results and model coefficients, are released without restriction alongside the analysis code. The published reference datasets used here, plasma proteomic signatures from the Global Neurodegeneration Proteomics Consortium and Mendelian randomization results from Yang et al., remain available from their original sources.

## Code availability

All analysis and figure-generation code is available at [repository DOI to be inserted on acceptance]. The repository reproduces every number, figure and supplementary panel reported here, and comprises the analysis pipeline, one script per main and supplementary figure, and verification scripts that recompute each quoted value from its source file and check figure panels against the text. A single configuration file holds every threshold used in the manuscript, so the reported criteria and the executed criteria cannot diverge. The runtime is recorded rather than asserted: R 4.4.1 (2024-06-14) with package versions captured at the time the figures were generated. Individual-level data are not included in the repository and are excluded by configuration; the analysis scripts expect them to be supplied separately under the access terms above.

## SUPPLEMENTAL FIGURE LEGENDS

**Supplementary Figure 1. Differential abundance and age-dependence for every disorder.**

Volcano plots for all 26 disorders with at least eight affected participants, one disorder per row. Left column: differential abundance against all available controls. Right column: the group x age interaction for the same disorder. Visual conventions follow Figure 2B, with the y axis showing nominal -log10(p) and the dashed line marking the FDR < 0.10 boundary. Panel A shows the 16 site-adjusted disorders and panel B the 10 site-confounded disorders; within each, disorders are ordered by descending number of differentially abundant proteins. WWOX-Related Disorder is not shown (with six affected participants it falls below the eight-participant threshold). The previous version of this legend referred to 27 disorders and to a pediatric-subset column; under multiple-testing correction the pediatric subset yields a single significant protein across all site-adjusted disorders, and it has been replaced by the interaction analysis.

**Supplementary Figure 2. Differential abundance calls are robust to the significance threshold but not to the model specification.**

Two distinct robustness questions, shown together because they have different answers and should not be conflated. (A, B) Effect of the significance criterion: per-disorder counts under three rules applied to the same model fits, namely the nominal p < 0.01 threshold used previously, the Benjamini-Hochberg FDR < 0.10 criterion used throughout this revision, and the CurveCurator hyperbolic boundary applied to FDR-adjusted p-values, which couples the fold-change and significance criteria rather than thresholding them independently. Disorder rank by perturbation burden is largely preserved across criteria in both the site-adjusted (Spearman rho = 0.894) and site-confounded (rho = 0.939) analyses. (C, D) Effect of the model specification: per-disorder counts at FDR < 0.10 under 1:1 matched controls as previously published, all available controls without a site term, and all controls with study site as a random effect, the model used throughout. Rank concordance is rho = 0.782 in the site-adjusted analysis but only rho = 0.406 in the site-confounded analysis, where disorder and site are collinear. (E, F) Number of proteins with an FDR-significant group x age interaction per disorder. The contrast between the rows is the point: how strictly significance is called has little effect on which disorders appear most perturbed, whereas how the data are modeled has a substantial effect. Robustness to the threshold should not be read as robustness to the model.

**Supplementary Figure 3. Robustness of the two-group solution.**

(A) Gap statistic and average silhouette width across k. The silhouette peaks at k = 2 (0.483 versus 0.364 at k = 3); the gap statistic rises monotonically, so k = 2 is selected by the one-standard-error rule rather than by a maximum. (B) Consensus clustering over 1,000 bootstrap resamples: mean within-group co-clustering probability 0.933 (minimum 0.567, SYNGAP1-Related Disorder). (C) Per-disorder silhouette width at k = 2; no disorder is negative. (D) Adjusted Rand index against the reported partition across four distance metrics and three linkages. Nine of twelve combinations reproduce it exactly; the three exceptions all use Euclidean distance, which tracks the number of differentially abundant proteins per disorder (r = 0.92) rather than profile similarity. (E) Position of each disorder along PC1, which separates the two clusters completely; site-adjusted and site-confounded disorders are distributed across the full range, so the split does not recover the analysis design.

**Supplementary Figure 4. Group-level enrichment and protein sharing across neurodegenerative signatures.**

(A) Group-level enrichment against each neurodegenerative signature. Per-disorder gene set enrichment p-values were combined within each cluster using Fisher’s method for combining independent p-values; tiles give the mean -log10 adjusted p across the disorders in each cluster, with the number of significant disorders and the combined p-value annotated. Note that this is Fisher’s combination METHOD applied to enrichment p-values and is unrelated to the Fisher exact overlap test used in the previous version of this analysis, which has been removed. (B) Overlap between the three protein sets used in this analysis. UpSet plot of proteins differentially abundant in at least one rare disorder in this study (5,836), proteins differentially abundant in at least one reference neurodegenerative signature (6,528), and proteins with colocalization-supported Mendelian randomization evidence (196), restricted to the 7,257 analytes shared across platforms. Bars give the size of each intersection; 5,152 proteins are shared between this study and the reference signatures, while 159 appear in all three.

**Supplementary Figure 5. Extended validation of the Proteomic Severity Index.**

(A) Vineland score selection. Number of severity-associated proteins at FDR < 0.05 for each of the five Vineland scores tested, from a global model fitted across all affected participants. Receptive Communication GSS yielded 223, against 59 for Expressive Communication and at most one for the Adaptive Behavior Composite, Communication Composite and Daily Living Skills, which is why Receptive GSS was chosen as the outcome for the multivariate model. (B) Association with Receptive versus Expressive Communication GSS across all 10,738 analytes, as −log10(p) from the global model. Points are colored by which outcome reaches FDR < 0.05: Receptive only, Expressive only, both, or neither; the dashed line is the identity. The ten most strongly Receptive-associated proteins that the index retains are labeled. Most signal lies below the identity line, indicating stronger association with Receptive GSS. (C) Per-disorder leave-one-disorder-out accuracy. Within-disorder Pearson correlation between observed and predicted Receptive GSS for each of the 22 disorders with at least five participants, colored by cluster (C1, C2). The dashed line marks the median of 0.55. Accuracy varies widely between disorders and is not systematically higher in either cluster. (D) Overlap between the 186 index proteins and four biomarker frameworks, tested by hypergeometric test against the full 10,738-analyte proteome. Proteins altered in neurodegenerative disease overlap the index more than expected by chance (125 observed against 113.6 expected, p = 0.05), though not after correction across the four tests (BH-adjusted p = 0.19). The Figure 4 convergent set (0 of 40), the Figure 5D convergent set (0 of 36) and the epilepsy meta-analysis set (0 of 32) share no protein with the index, so the severity proteins are a distinct set rather than a subset of the cross-disorder biomarkers. (E) Expression of the 40 index proteins with the largest absolute coefficients across the disorders with Vineland data, grouped by cluster. Color is the index regression coefficient per standard deviation of Receptive GSS, not a case-control fold change; dot size gives the within-disorder association p-value.

**Supplementary Figure 6. Recurrence below the cross-disorder convergence threshold.**

(A) Number of analytes reaching FDR < 0.10 in one, two, or three or more of the 16 site-adjusted disorders. Of 1,992 analytes significant in at least one disorder, 1,571 are significant in only one and 369 in exactly two; 52 reach three or more, of which 40 also satisfy the 70% directional-consistency requirement and constitute the convergent set of Figure 4. (B) Direction of change irrespective of significance. For every analyte measured in all 16 site-adjusted disorders, the number of disorders sharing the dominant sign of log2 fold-change. Proteins named in the Discussion are marked. ATF6, COMP and AKR1C3 shift consistently in 15 of 16 disorders and GDF15, CCL11, HSD11B1 and MOB4 in 14 of 16, but 688 analytes reach 14 of 16 and 212 reach 15 of 16, so a skew of this size indicates a shared trend rather than exceptional convergence. (C) Per-disorder support for the proteins named in the Discussion, shown separately for the site-adjusted and site-confounded analyses and colored by direction. The dashed line marks the three-disorder convergence threshold, applied within each analysis. Only ATF6 and COMP meet it in the site-adjusted analysis; GDF15, CCL11 and HSD11B1 fall below it, and MOB4 has no site-adjusted support. CXCL3, C3 and MOB4 change in both directions across the site-confounded disorders, which is why they are not treated as directionally convergent.

**Supplementary Figure 7. The severity index against time since last seizure.**

Time since the last seizure was recorded at collection for 257 of 522 affected participants; 222 of these have confirmed epilepsy, an index value and an age, and form the analysis set. (A) Proteomic Severity Index against time since last seizure on a logarithmic scale, colored by cluster, with a linear fit and 95% confidence band. Higher index values, indicating better receptive communication, occur at longer intervals since the last seizure (r = 0.32; 0.29 after adjustment for age). (B) The same correlation computed within each of the 12 disorders contributing at least eight participants. It is positive in nine, with a median of 0.17, so the association is not solely between disorders, though it is substantially weaker within them. (C) Null distribution from 2,000 permutations of seizure recency within disorder, which holds the between-disorder structure fixed. The null is centered at 0.19 rather than zero, confirming that most of the raw association reflects differences between disorders; the observed age-adjusted correlation of 0.29 exceeds it at p = 0.012. Time since last seizure is a single retrospective field that conflates seizure control with seizure severity and is missing for roughly half the affected cohort, so this analysis supports the interpretive caution stated in the Discussion rather than establishing a quantitative relationship between seizure burden and the index.

