## Supplemental Figures for "A plasma proteomic atlas of rare genetic neurodevelopmental disorders reveals convergent molecular signatures, neurodegenerative overlap, and a cross-disorder severity index"

### Differential abundance and age−dependence for every disorder

#### Tier 1 − site−adjusted (16 disorders)

**Differential expression (all ages)** | **Group x age interaction**

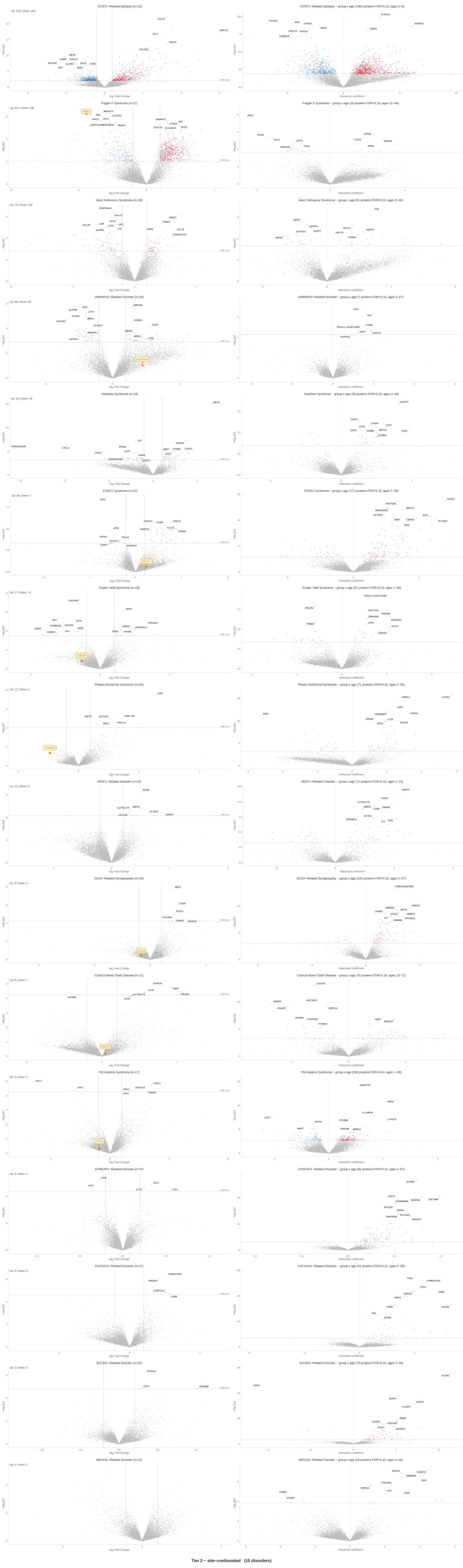


#### Tier 2 − site−confounded (10 disorders)

**Differential expression (all ages)** | **Group x age interaction**

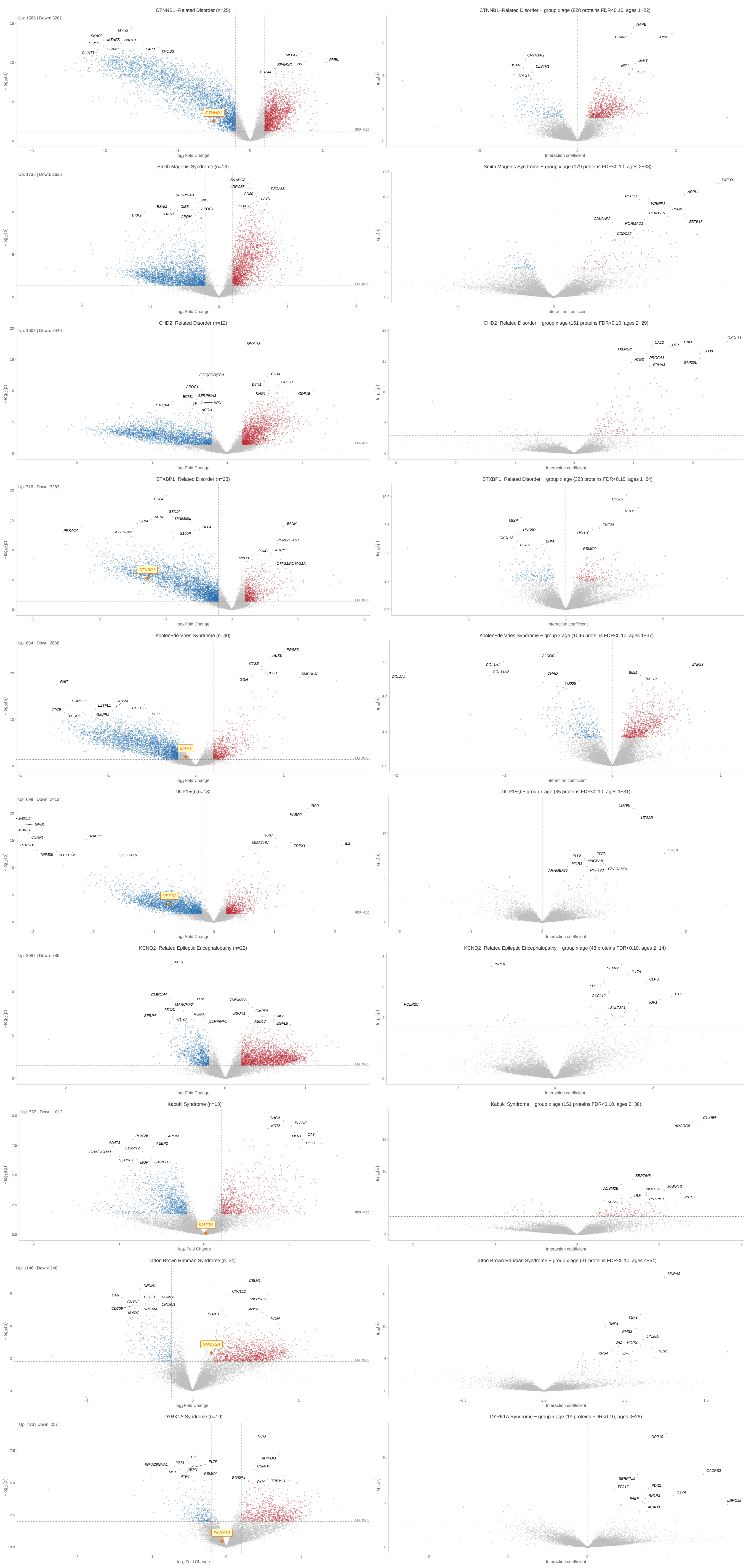

#### DEP calls are robust to the significance threshold but not to the model

A, B: three significance criteria applied to the same fits. C, D: three model specifications at FDR < 0.1
E, F: proteins whose disease effect varies with age, per disorder.
Disorder rank is preserved across thresholds but not across models − the modelling choice matters more than the cutoff.

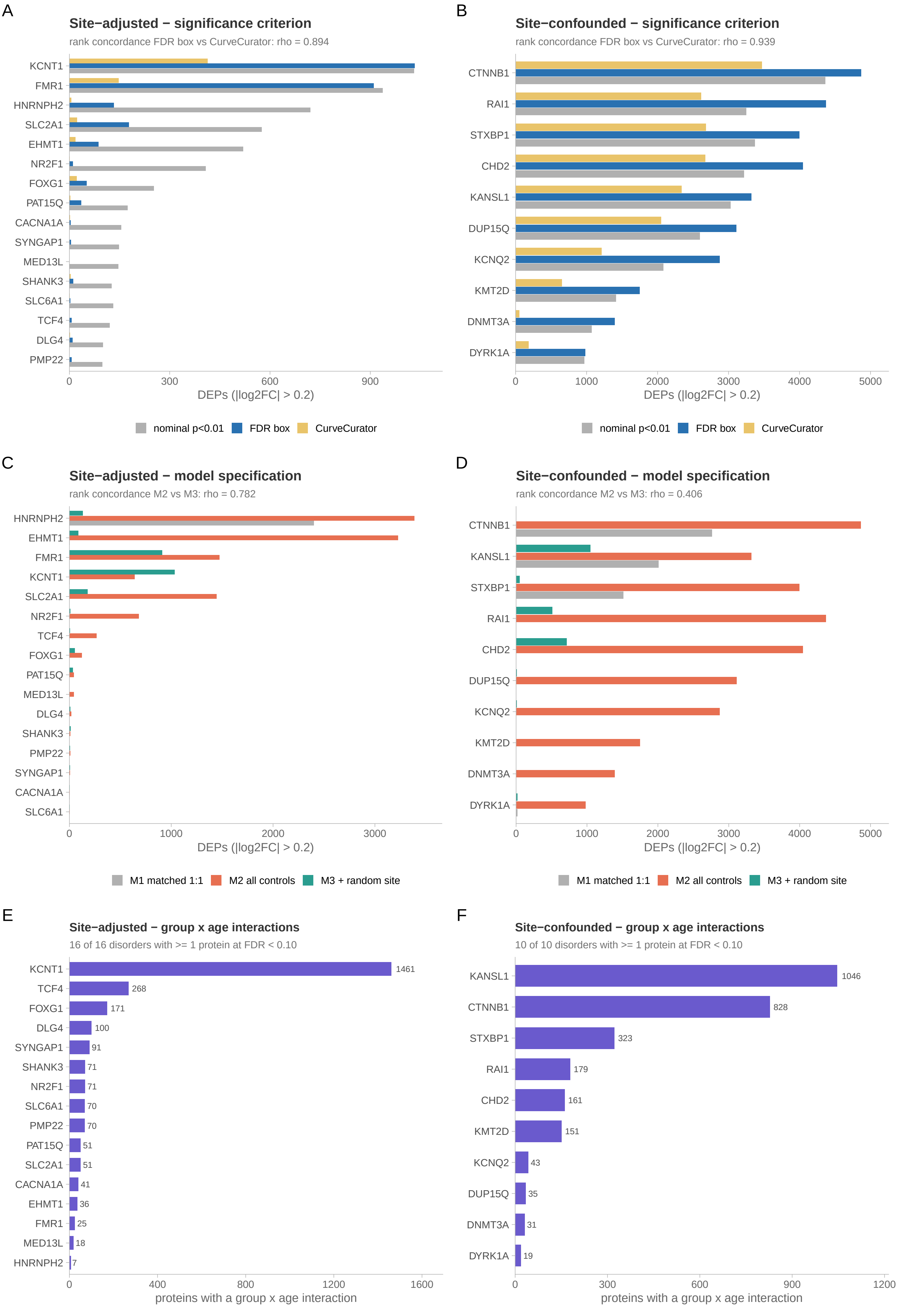

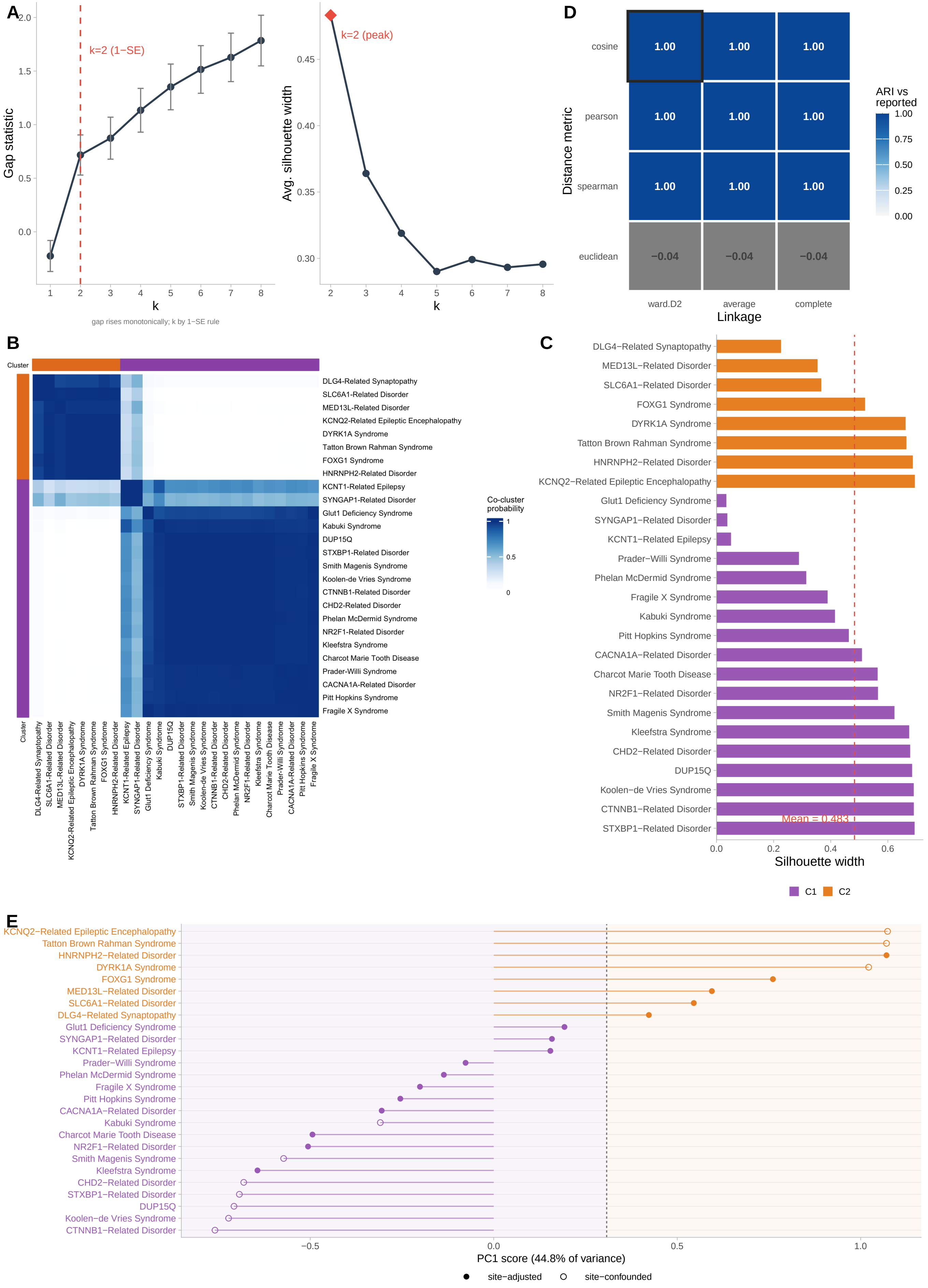

A
Gap statistic
k=2 (1−SE)
k
gap rises monotonically; k by 1−SE rule
Avg. silhouette width
k=2 (peak)
k
D
Distance metric
cosine
pearson
spearman
euclidean
1.00
−0.04
ward.D2
average
complete
Linkage
ARI vs reported
B
Cluster
DLG4-Related Synaptopathy
SLC6A1-Related Disorder
MED13L-Related Disorder
KCNQ2-Related Epileptic Encephalopathy
DYRK1A Syndrome
Tatton Brown Rahman Syndrome
FOXG1 Syndrome
HNRNPH2-Related Disorder
KCNT1-Related Epilepsy
SYNGAP1-Related Disorder
Glut1 Deficiency Syndrome
Kabuki Syndrome
DUP15Q
STXBP1-Related Disorder
Smith Magenis Syndrome
Koolen-de Vries Syndrome
CTNNB1-Related Disorder
CHD2-Related Disorder
Phelan McDermid Syndrome
NR2F1-Related Disorder
Kleefstra Syndrome
Charcot Marie Tooth Disease
Prader-Willi Syndrome
CACNA1A-Related Disorder
Pitt Hopkins Syndrome
Fragile X Syndrome
Co-cluster probability
C
Mean = 0.483
Silhouette width
C1
C2
E
PC1 score (44.8% of variance)
site−adjusted
site−confounded

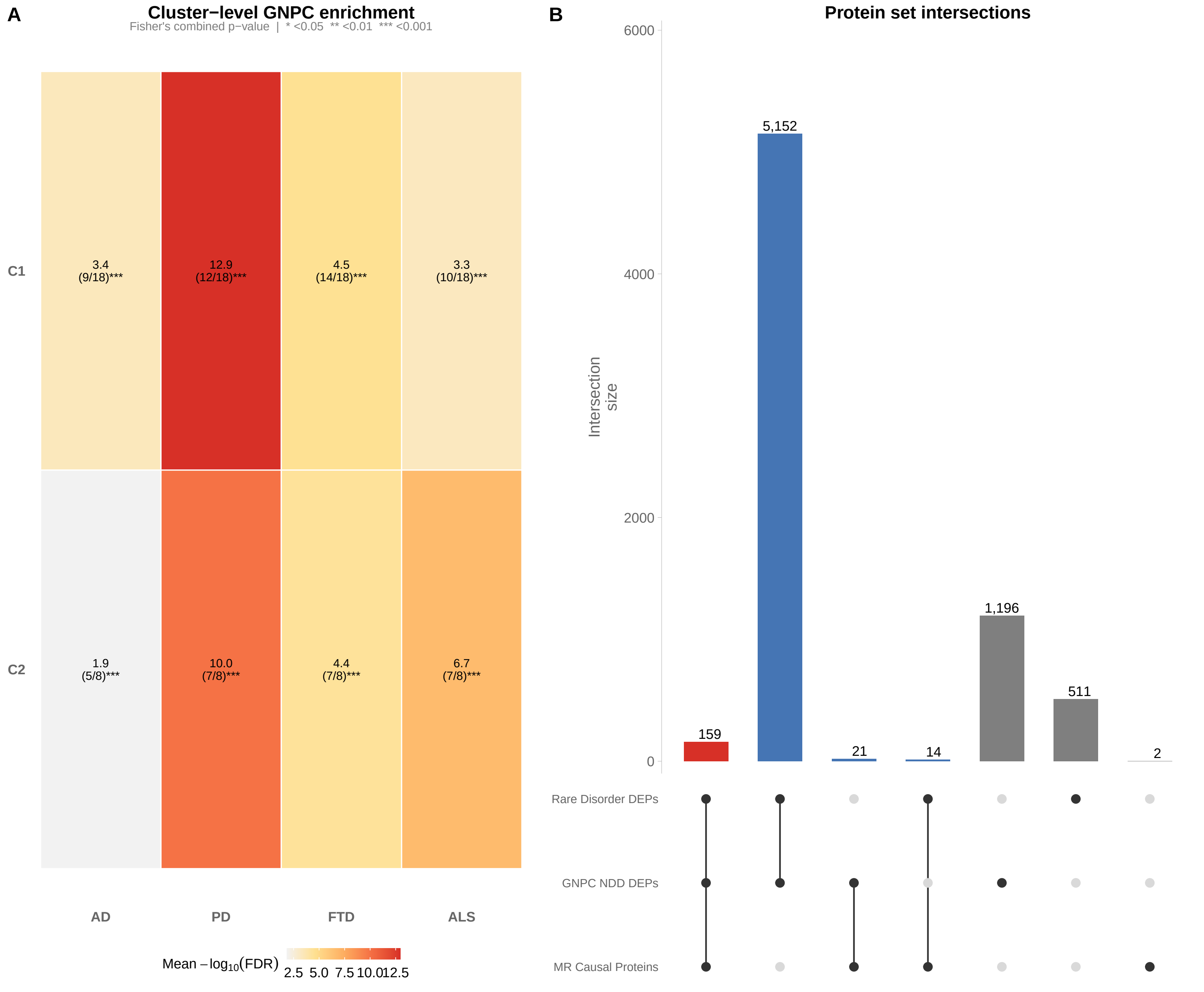
A
Cluster−level GNPC enrichment
Fisher's combined p−value | * <0.05 ** <0.01 *** <0.001
C1
3.4 (9/18)***
12.9 (12/18)***
4.5 (14/18)***
3.3 (10/18)***
C2
1.9 (5/8)***
10.0 (7/8)***
4.4 (7/8)***
6.7 (7/8)***
AD
PD
FTD
ALS
Mean −log10(FDR)
2.5 5.0 7.5 10.0 12.5
B
Protein set intersections
Intersection size
6000
4000
2000
0
159
5,152
21
14
1,196
511
2
Rare Disorder DEPs
GNPC NDD DEPs
MR Causal Proteins

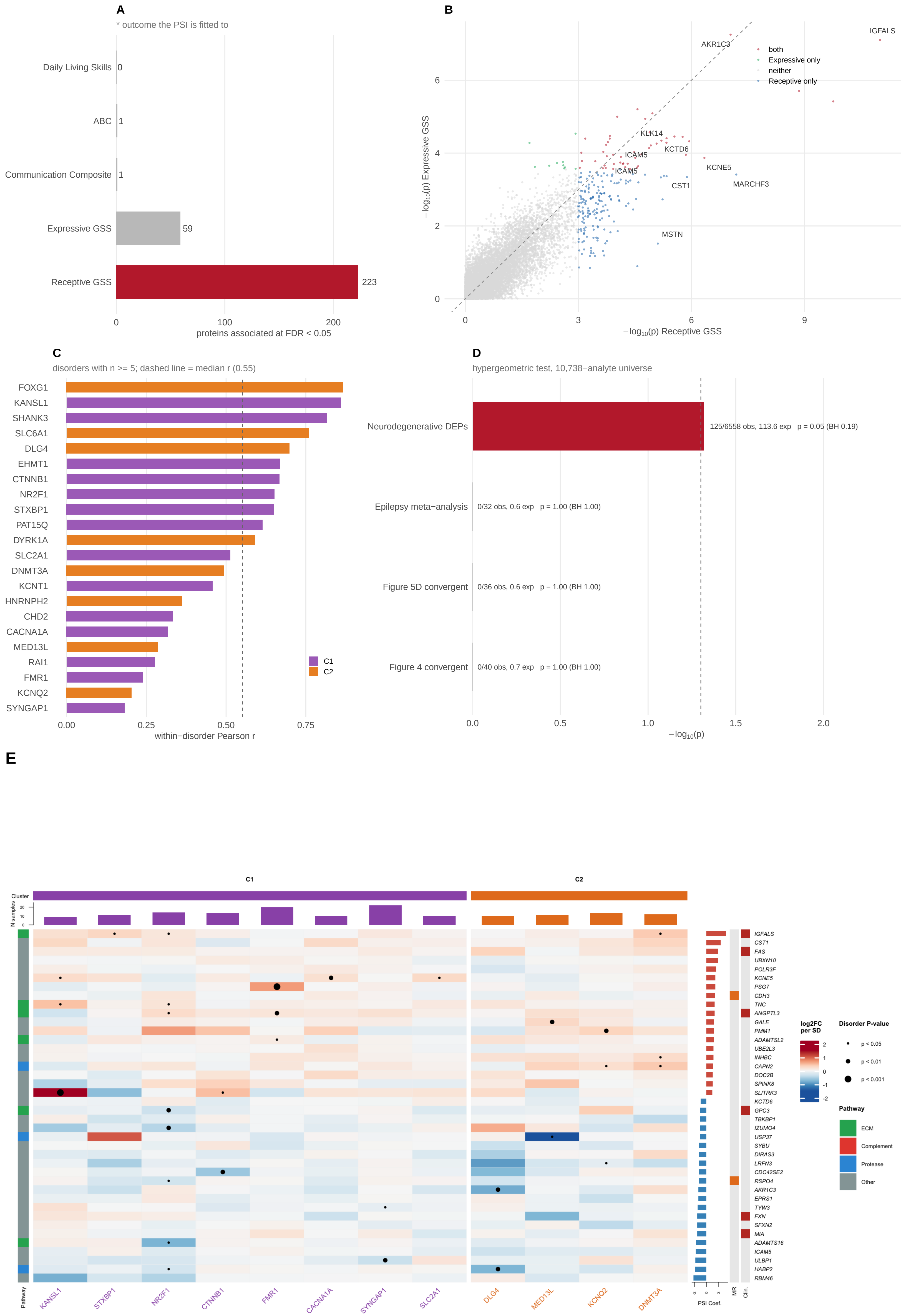

A
* outcome the PSI is fitted to
Daily Living Skills 0
ABC 1
Communication Composite 1
Expressive GSS 59
Receptive GSS 223
0
100
200
proteins associated at FDR < 0.05
B
both
Expressive only
neither
Receptive only
IGFALS
AKR1C3
KLK14
KCTD6
ICAM5
ICAM5
KCNE5
CST1
MARCHF3
MSTN
−log10(p) Expressive GSS
−log10(p) Receptive GSS
0
2
4
6
0
3
6
9
C
disorders with n >= 5; dashed line = median r (0.55)
FOXG1
KANSL1
SHANK3
SLC6A1
DLG4
EHMT1
CTNNB1
NR2F1
STXBP1
PAT15Q
DYRK1A
SLC2A1
DNMT3A
KCNT1
HNRNPH2
CHD2
CACNA1A
MED13L
RAI1
FMR1
KCNQ2
SYNGAP1
C1
C2
0.00
0.25
0.50
0.75
within−disorder Pearson r
D
hypergeometric test, 10,738−analyte universe
Neurodegenerative DEPs
125/6558 obs, 113.6 exp p = 0.05 (BH 0.19)
Epilepsy meta−analysis
0/32 obs, 0.6 exp p = 1.00 (BH 1.00)
Figure 5D convergent
0/36 obs, 0.6 exp p = 1.00 (BH 1.00)
Figure 4 convergent
0/40 obs, 0.7 exp p = 1.00 (BH 1.00)
0.0
0.5
1.0
1.5
2.0
−log10(p)
E
C1
C2
Cluster
N samples
20
10
0
IGFALS
CST1
FAS
UBXN10
POLR3F
KCNE5
PSG7
CDH3
TNC
ANGPTL3
GALE
PMM1
ADAMTSL2
UBE2L3
INHBC
CAPN2
DOC2B
SPINK8
SLITRK3
KCTD6
GPC3
TBKBP1
IZUMO4
USP37
SYBU
DIRAS3
LRFN3
CDC42SE2
RSPO4
AKR1C3
EPRS1
TYW3
FXN
SFXN2
MIA
ADAMTS16
ICAM5
ULBP1
HABP2
RBM46
Pathway
KANSL1
STXBP1
NR2F1
CTNNB1
FMR1
CACNA1A
SYNGAP1
SLC2A1
DLG4
MED13L
KCNQ2
DNMT3A
PSI Coef.
MR
Clin.
log2FC per SD
2
1
0
-1
-2
Disorder P-value
p < 0.05
p < 0.01
p < 0.001
Pathway
ECM
Complement
Protease
Other

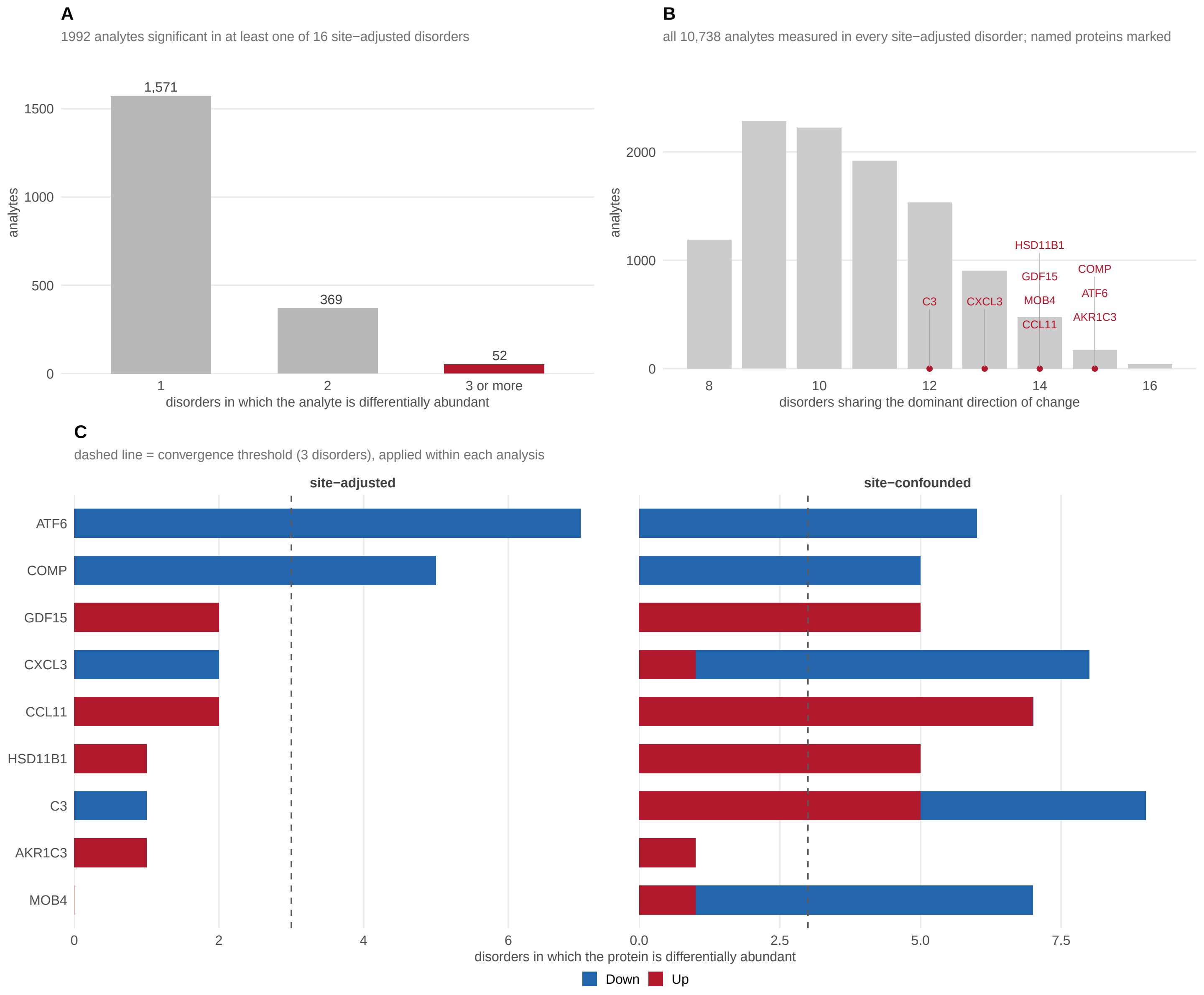
A
1992 analytes significant in at least one of 16 site−adjusted disorders
1,571
369
52
1500
1000
500
0
analytes
1
2
3 or more
disorders in which the analyte is differentially abundant
B
all 10,738 analytes measured in every site−adjusted disorder; named proteins marked
2000
1000
0
analytes
HSD11B1
COMP
GDF15
ATF6
C3
CXCL3
MOB4
AKR1C3
CCL11
8
10
12
14
16
disorders sharing the dominant direction of change
C
dashed line = convergence threshold (3 disorders), applied within each analysis
site−adjusted
site−confounded
ATF6
COMP
GDF15
CXCL3
CCL11
HSD11B1
C3
AKR1C3
MOB4
0
2
4
6
0.0
2.5
5.0
7.5
disorders in which the protein is differentially abundant
Down
Up

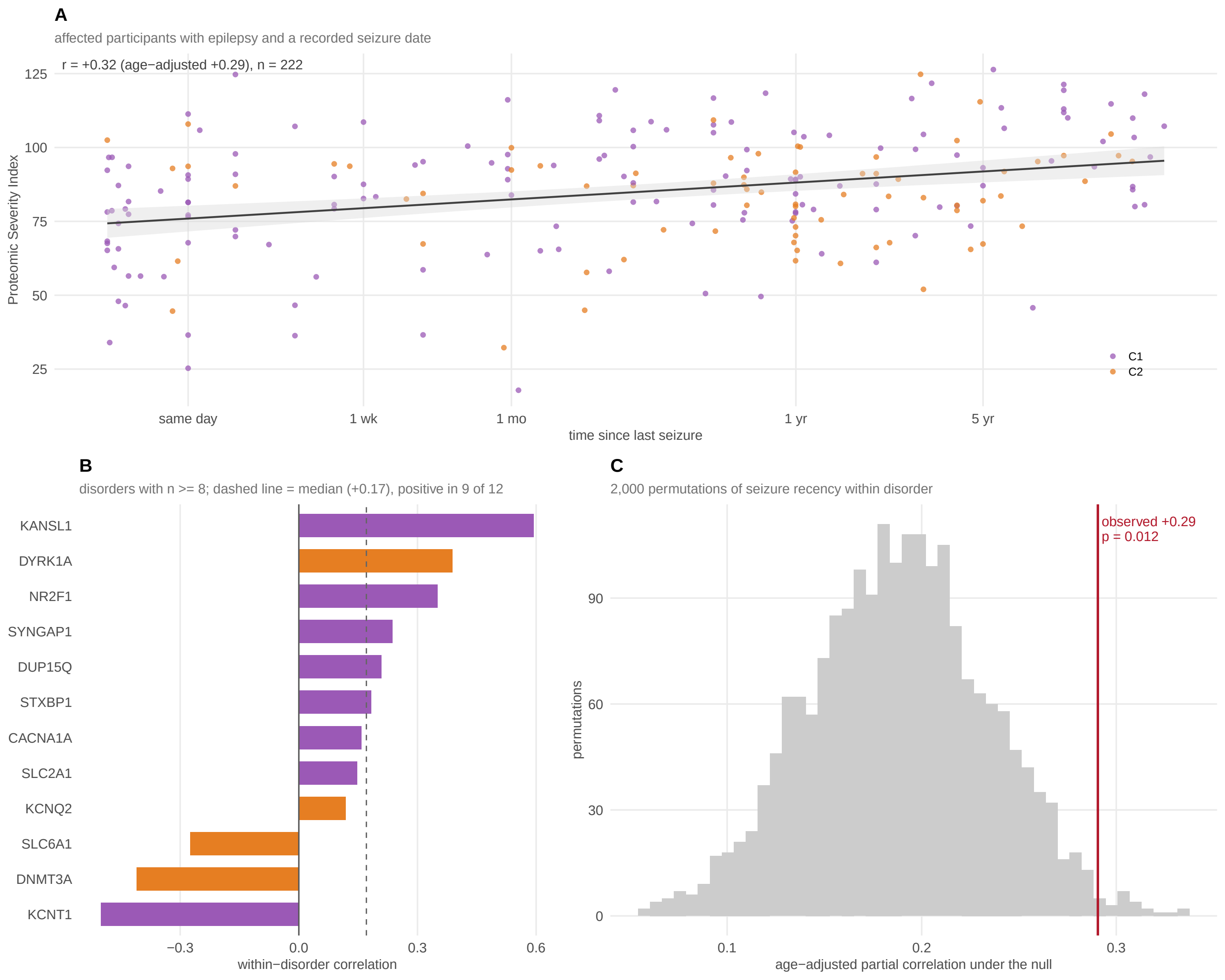
A
affected participants with epilepsy and a recorded seizure date
r = +0.32 (age−adjusted +0.29), n = 222
Proteomic Severity Index
125
100
75
50
25
same day
1 wk
1 mo
1 yr
5 yr
time since last seizure
C1
C2
B
disorders with n >= 8; dashed line = median (+0.17), positive in 9 of 12
KANSL1
DYRK1A
NR2F1
SYNGAP1
DUP15Q
STXBP1
CACNA1A
SLC2A1
KCNQ2
SLC6A1
DNMT3A
KCNT1
−0.3
0.0
0.3
0.6
within−disorder correlation
C
2,000 permutations of seizure recency within disorder
observed +0.29
p = 0.012
permutations
90
60
30
0
0.1
0.2
0.3
age−adjusted partial correlation under the null
